# Genotype-dependent reductions of competition are associated with yield maintenance in a modern common wheat *Triticum aestivum* L. cultivar

**DOI:** 10.64898/2026.09.07.749990

**Authors:** Nao Kosugi, Takuto Kaneko, Kentaro Yoshida, Akira Yamawo

**Affiliations:** Department of Biology, Faculty of Agriculture and Life Science, Hirosaki University, Hirosaki, Aomori 036-8560, Japan; Graduate School of Science, Kyoto University, Kitashirakawa Oiwake-cho, Sakyo-ku, Kyoto 606-8502, Japan; Center for Ecological Research, Kyoto University, Otsu, Shiga 520-2113, Japan; Graduate School of Agriculture, Kyoto University, Kitashirakawa Oiwake-cho, Sakyo-ku, Kyoto 606-8502, Japan

**Keywords:** genotype discrimination, conditional competitiveness, crop breeding, monoculture, root exudates

## Abstract

Modern crop breeding has increased yield under monoculture stands, but the mechanisms underlying this improvement remain incompletely understood. We tested whether genotype-dependent reduction of competition, whereby plants adjust competitive responses according to neighbor genetic identity, contributes to yield maintenance in modern common wheat cultivars. We conducted a root-exudate application experiment and two competition experiments using three common wheat (*Triticum aestivum* L.) landraces and one modern cultivar, Norin 61. Norin 61 exhibited little change in shoot-to-root allocation in response to root exudates from the same cultivar compared with distilled water, whereas it increased shoot-to-root allocation exposed to root exudate from different cultivars. Moreover, shoot-to-root allocation increased with decreasing genetic relatedness between neighboring cultivars in common wheat plants. In competition experiments, greater shoot-to-root allocation was associated with stronger suppression of competitors. Norin 61 maintained grain biomass under the same-cultivar competition compared to solitary conditions but exhibited substantial yield reductions when competing with different cultivars, whereas these patterns were weaker or absent in landraces. These results suggest that genotype-dependent reduction of competition may contribute to yield maintenance in modern wheat cultivars and provide a new perspective on how crop breeding has shaped plant–plant interactions in monoculture stands.

**Highlight:** Genotype discrimination may reduce intraspecific competition among genetically identical wheat plants, helping modern cultivar maintain yield in monoculture stands.

## Introduction

Plants frequently compete with neighboring individuals for limiting resources such as light, water, and soil nutrients. Under belowground competition, plants often allocate additional resources to root growth to enhance nutrient acquisition, resulting in reduced reproductive investment (Gersani *et al*., 2001; Maina *et al*., 2002; O’Brien *et al*., 2005). This phenomenon, known as the “tragedy of the commons”(Gersani *et al*., 2001), arises when selection favors competitive traits that increase individual resource acquisition but reduce the overall productivity of the population (Kiers and Denison, 2014). Such overinvestment in competitive structures is expected to decrease crop yields, particularly under dense monoculture stands where neighboring individuals compete for the same resources (Chen *et al*., 2012; Kiers and Denison, 2014).

Modern crop breeding has substantially increased yield under monoculture stands despite this evolutionary conflict. Classical crop ideotype theory proposes that artificial selection has achieved this improvement by favoring individuals with reduced competitive ability and greater allocation to yield production, thereby mitigating “the tragedy of the commons” within crop populations (Donald, 1968; Sedgley, 1991; Weiner, 2003). However, this explanation assumes that competitiveness is uniformly reduced. Under agricultural conditions, plants frequently encounter genetically identical neighbors belonging to the same cultivar, but they may also encounter genetically distinct cultivars or heterospecific competitors such as weeds. Consequently, complete loss of competitive ability may not always be advantageous. An alternative possibility is that modern cultivars express competitive traits conditionally, suppressing competition when surrounded by genetically similar individuals while maintaining the capacity to compete when interacting with genetically distinct neighbors. Such conditional competitiveness could simultaneously maximize stand-level productivity and preserve the ability to acquire resources in heterogeneous competitive environments.

One mechanism that may enable conditional competitiveness is genotype discrimination, whereby plants modify their growth and resource allocation according to the genetic identity of neighboring conspecifics (Chen *et al*., 2012; Murphy *et al*., 2017*a*; Anten and Chen, 2021; Sher *et al*., 2025; Yamawo, 2026). Previous studies have demonstrated that many plant species alter biomass allocation, morphology, and competitive behavior depending on the genetic relatedness of neighboring individuals, genetic phenotype-dependent response, which is referred to as kin discrimination in wild plants (Dudley and File, 2007; Murphy and Dudley, 2007; Fang *et al*., 2011; Crepy and Casal, 2015; Yamawo, 2015, 2021; Yamawo *et al*., 2017; Yang *et al*., 2018; Fukano *et al*., 2019; Takigahira and Yamawo, 2019; Yamawo and Mukai, 2020). Kin discrimination is interpreted as the result that plants distinguish genetically related individuals from genetically unrelated ones, referring to kin recognition (Anten and Chen, 2021; Mazal *et al*., 2023). In many cases, plants exhibit reduced investment in competitive traits when interacting with genetically similar individuals but increase competitive allocation when exposed to genetically distinct neighbors (Yamawo, 2026). Such plastic responses are thought to reduce unnecessary competition among genetically related individuals while maintaining the capacity to compete when resource acquisition is threatened (Novoplansky, 2019; Anten and Chen, 2021).

Similar genotype-dependent responses, genotype discrimination, have also been reported in crop species, where they may influence competition among cultivars or genotypes (Ninkovic, 2003; Fang *et al*., 2013; Zhu and Zhang, 2013; Murphy *et al*., 2017*b*; Yang *et al*., 2018; Saleh and Kniss, 2022). These studies suggest that discrimination among neighboring genotypes may influence competitive interactions and potentially improve crop performance under dense planting conditions. At the same time, several reports have argued that genotype discrimination resulting from its recognition should decline during domestication and breeding because artificial selection favors reduced competitiveness and greater uniformity among individuals (Kiers and Denison, 2014; Fukano *et al*., 2019). This creates an unresolved paradox. If modern crop breeding favors reduced competition among neighboring plants, why do genotype-dependent responses persist in many cultivated species? One possibility is that breeding has not eliminated competitive ability itself but has instead favored mechanisms that allow plants to deploy competitive traits selectively according to neighbor identity. However, empirical evidence directly linking genotype discrimination to yield maintenance in modern crop cultivars remains limited.

Common wheat (*Triticum aestivum* L.) provides an ideal system for addressing this question because it includes both landraces and modern cultivars that differ substantially in breeding history, genetic background, and productivity under monoculture stands (Takenaka *et al*., 2018). Previous work using one landrace and one modern cultivar has shown that the modern common wheat cultivar can alter its growth in response to the genotype of neighboring plants, but the landrace did not show such response (Zhu and Zhang, 2013). However, it remains largely unclear whether modern cultivars suppress competitive responses toward individuals of the same cultivar by discrimination of genetic identity, whether such responses differ from those of landraces, and whether they are associated with yield maintenance under competitive conditions.

In this study, we examined the possibility that modern wheat breeding may favor genotype-dependent reduction of competition among individuals of the same cultivar. Using Norin 61 as a focal modern cultivar and three landraces, we tested whether competitive allocation varies according to neighbor genotypes and whether such variation is associated with yield maintenance under same-cultivar competition. To address these questions, we conducted three complementary experiments. First, we examined whether variation in shoot-to-root allocation, one of the most commonly used indices of kin or genotype discrimination (Dudley *et al*., 2013), represents a competitive trait by testing its effects on neighboring plants. If increased shoot-to-root allocation enhances competitive ability, individuals expressing this trait should suppress the growth of their competitors. Second, we investigated genotype-dependent responses using root exudates, which are known to act as cues for neighbor discrimination (Biedrzycki et al., 2010; Semchenko et al., 2014; Yamawo et al., 2017; Yang et al., 2018). We predicted that Norin 61 would suppress competitive allocation when exposed to root exudates from the same cultivar but retain competitive responses when exposed to exudates from genetically distinct cultivars. We then compared this response with those of the three landraces to assess variation among cultivars. Finally, we compared yield performance under solitary, same-cultivar, and different-cultivar competition. If genotype-dependent reduction contributes to yield maintenance, cultivars that reduce competitive allocation toward individuals of the same cultivar should experience smaller reductions in yield under same-cultivar competition.

## Materials and methods

### Plant materials

We used four common wheat (*Triticum aestivum* L.) cultivars: Norin 61 (N61), Nobeokabouzu (NBB), Chinese Spring (CS), and KU-7113 (NP) (Table 1). N61 is a modern cultivar, whereas the others are landraces. Population structure analysis demonstrated that N61 and NBB were classified in the same cluster, and CS and NP were in the same cluster, and N61, NBB, CS, and NP were genetically distant in that order (Takenaka *et al*., 2018). All seeds were obtained from the National BioResource Project Wheat, Japan. *Triticum turgidum* L. subsp. *durum* (Desf.) Husnot cv. ‘Langdon’ was additionally included as a distantly related competitor, heterospecific competitor, in mixed-cultivar treatments.

**Table 1.**
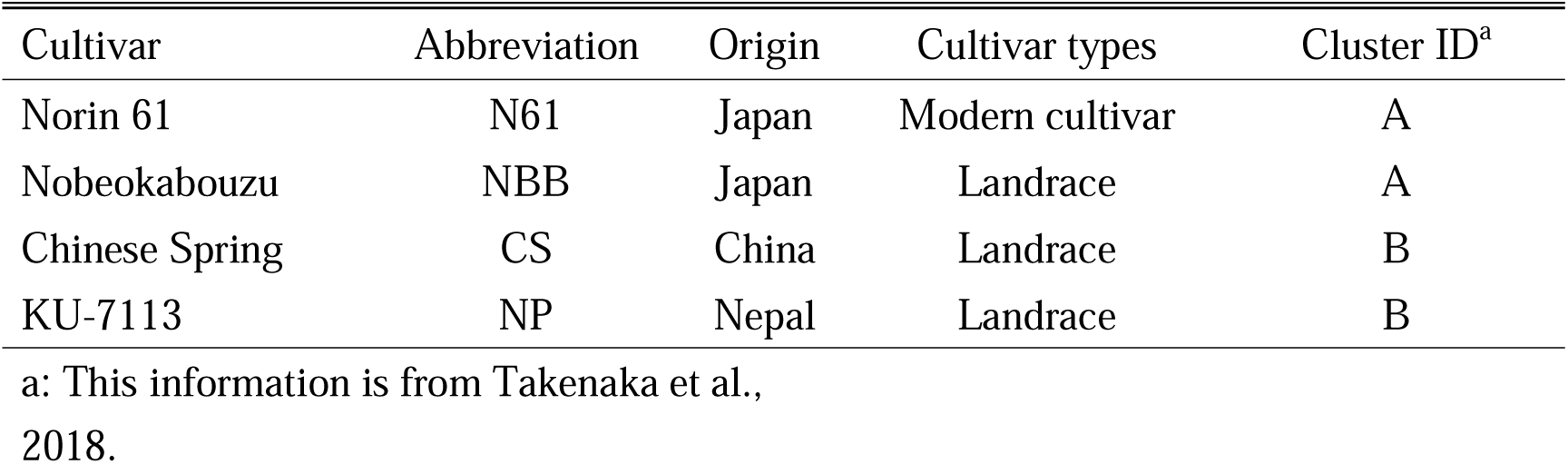
Summary information about four common wheat cultivars from National BioResource Project (NBRP) Wheat.

| Cultivar | Abbreviation | Origin | Cultivar types | Cluster ID<sup>a</sup> |
| --- | --- | --- | --- | --- |
| Norin 61 | N61 | Japan | Modern cultivar | A |
| Nobeokabouzu | NBB | Japan | Landrace | A |
| Chinese Spring | CS | China | Landrace | B |
| KU-7113 | NP | Nepal | Landrace | B |

a: This information is from Takenaka et al., 2018.

### Exp. 1: Competition experiment

Because our hypothesis predicts that modern cultivars suppress competitive allocation toward genetically identical neighbors, we first examined whether variation in shoot-to-root allocation is associated with competitive ability in common wheat plants. We conducted a competition experiment to test whether individuals with higher shoot-to-root ratios suppress the growth of neighboring plants. On June 21, 2022, seeds of each cultivar were germinated in Petri dishes containing moistened filter paper. The seeds were stratified at 4 °C for two days and then grown in a chamber at 25 °C under a 12-h light:12-h dark photoperiod for five days. On June 28, 2022, seedlings were randomly selected and transplanted into experimental pots (15 cm in diameter, 12.5 cm in height, and 1.6 L in volume) containing a mixture of 70% horticultural soil (Nippi Horticultural Soil No. 1; Nihon Hiryo Co., Ltd.) and 30% red soil.

We established two treatments: pairs of the same cultivar (30 pairs) and pairs of different cultivars (24 pairs) (Table S1). In the paired treatments, two seedlings were planted 5 cm apart in each pot. Different-cultivar pairs included combinations involving *Triticum turgidum* L. subsp. *durum* (Desf.) Husnot cv. ‘Langdon’ as a distantly related, heterospecific competitor. All pots were maintained in a greenhouse at Hirosaki University (40°58′ N, 140°47′ E) for 38 days and watered every day. At the end of the experiment, all individuals were harvested and dried at 60 °C for one week. The shoot and root biomass of each individual were measured separately.

### Exp.2: Root-exudation application

#### Root exudate collection

To determine whether common wheat cultivars adjust their competitive allocation according to neighbor genotypes, we conducted a root-exudate application experiment. The root exudates of each cultivar were collected using a hydroponic cultivation method based on a previously established protocol for studying kin or genotype discrimination in rice (Fig.S1) (Yang *et al*., 2018). On January 16, 2023, seeds of each cultivar were germinated in Petri dishes containing moistened filter paper. Seeds were stratified at 4 for two days. After that, they were grown in a chamber at 25 under a 12-h light:12-h dark photoperiod for 5 days. Twenty-five seedlings of each cultivar as emitter plants were inserted into holes in a Styrofoam plate with twenty-five holes (13 cm length, 10.5 cm width). We transplanted two Styrofoam plates with seedlings into a plastic container (21.5 cm length, 15.2 cm width) containing 300 mL of distilled water (Fig.S1 a). Root exudates from each cultivar were collected, just prior to application. This process was performed every day during the experimental period. After root exudate collection, the same amount of distilled water was added to each plastic container. Each collected root exudate was used in the root exudate application experiment described below.

#### Root exudate application

Seedlings of each cultivar as receiver plants was transplanted into a 10 mL pipette tip containing silica sand (Fig.S1 b). Seedlings from each cultivar were assessed in each of the three experimental treatments: (1) 4 mL of distilled water in a 10 mL pipette tip (24 individuals); (2) 4 mL of the same cultivar’s root exudate in a 10 mL pipette tip (24 individuals); and (3) 4 mL of different cultivars’ root exudate in a 10 mL pipette tip (24 seedlings for each different cultivar) (Fig.S1 b; Table S2). For each receiver cultivar, root exudates from each of the other three cultivars were applied separately rather than pooled, allowing us to evaluate responses to individual donor cultivars. Distilled water and root exudates from the same or different cultivars were applied every day and immediately after root exudate collection. All seedlings were grown in a growth chamber at 25 under a 12-h light:12-h dark photoperiod for 15 days. All individuals were collected and dried at 70 °C for one week. We measured the shoots and roots of each individual separately.

### Exp.3: Yield measurements

To evaluate whether genotype discrimination resulting from its recognition contributes to yield maintenance under monoculture stands, we compared yield performance among solitary, same-cultivar, and different-cultivar competition treatments. Seeds of each cultivar were germinated in Petri dishes with moistened filter paper in October 2022. Seeds were stratified at 4 for two days. After that, they were grown in a chamber at 25 □ under a 12-h light:12-h dark photoperiod for 5 days. We randomly selected and transplanted seedlings into the experimental pots (15 cm in diameter, 12.5 cm in height, and 1.6 L in volume) containing 70 % horticultural soil (Nippi Horticultural Soil No.1) (Nihon Hiryo Co., Ltd.) and 30 % red soil.

We used three treatments: solitary (30 individuals), pairs of the same cultivar (30 pairs), and pairs of different cultivars (32 pairs) (Table S3). In addition, the different-cultivar treatment included *T. turgidum* subsp*. durum* cv. ‘Langdon’ as a distantly related competitor, which is a different species. The distance between the paired seedlings was 5 cm. Each of the 30 pots with a single seedling was divided in half using a plastic plate, and one seedling was planted in one half of the pot. The other half of the pots were filled with mixed soil but had no plants. All pots were placed in a greenhouse at Hirosaki University (40 °58′ N, 140 °47′ E) for 40 days, then placed outdoors until harvest and watered every day. All individuals were harvested 50 days after heading and dried at 70 °C for one week.

We measured major yield-related traits, including panicles per plant, total grains per panicle, grain number, grain biomass, thousand-grain weight, and total aboveground biomass. Harvest index is seed biomass divided by total aboveground biomass. We also estimated field yield from grain biomass and pot surface area and expressed it as kg/ha.

### Data analysis

All statistical analyses were performed using R version 4.4.1 (R Development Core Team).

To evaluate whether shoot-to-root allocation represents a competitive trait in common wheat plants, we analyzed the relationship between shoot-to-root ratio and the biomass of neighboring individuals using linear mixed-effects models (LMMs). Shoot-to-root ratio of focal plant was included as a fixed effect, whereas cultivar identity and competition treatment were included as random effects. We also included all competition treatments, including those involving *Triticum turgidum* L. subsp. *Durum* (Desf.) Husnot cv. ‘Langdon’. In this analysis, we used both individuals planted as pairs, but we did not treat *T. turgidum* L. subsp. *durum* (Desf.) Husnot cv. ‘Langdon’ as focal plants. The significance of fixed effects was assessed using *F*-test.

To evaluate cultivar responses to root exudates, we first tested the effects of treatment on total biomass separately for each cultivar using generalized linear model (GLM) with a Gaussian distribution and identity link. Treatments consisted of distilled water, same-cultivar root exudates, and different-cultivar root exudates. To investigate differences in biomass allocation between shoots and roots, we analyzed shoot biomass as a function of root biomass, treatment, and their interaction using GLM with a Gaussian distribution and an identity link. When the interaction term was not significant, it was removed and the model was refitted with root biomass and treatment as explanatory variables. For analyses involving different-cultivar exudates, responses to each donor cultivar were treated separately. The significance of the explanatory variables was assessed by *F*-test. *P*-values were adjusted for multiple comparisons using the false discovery rate (FDR) method.

To evaluate whether genotype discrimination contributes to yield maintenance under competitive conditions, we analyzed harvest index and yield-related traits, including panicles per plant, grains per panicle, grain number, thousand-grain weight, total aboveground biomass, grain biomass, and estimated field yield. For each trait except for harvest index, panicle per plant and grain number, we tested the effects of cultivar, competition treatment (solitary, same-cultivar pair, and different-cultivar pair), and their interaction using GLM with a Tweedie distribution and log link by χ^2^ - test, because the data were overdispersal. Harvest index was analyzed using a GLM with a Gaussian distribution and identity link by *F*-test, because there was no overdispersal. Panicle per plant and grain number were analyzed using a GLM with a zero-inflated Poisson (ZIP) distribution and log link by χ^2^ - test, because the data were overdispersal. In these analyses, competition treatments involving *T. turgidum* L. subsp. *durum* (Desf.) Husnot cv. ‘Langdon’ were included in the different-cultivar category.

Individuals that died before harvest were assigned a value of zero for yield-related traits. However, in paired treatments, surviving individuals whose partner had died were excluded because their competitive environment differed from that of the remaining paired plants. *P*-values were adjusted for multiple comparisons using the FDR method.

## Results

### Exp.1: Competition experiment

Shoot-to-root allocation was negatively associated with the total biomass of neighboring individuals (estimated coefficient = −186.067, d.f. = 319.766, *F* = 48.119, *P* < 0.001; Fig. 1). Individuals with higher shoot-to-root ratios suppressed competitor growth more strongly, regardless of whether competitors belonged to the same or different cultivar or species (Fig. S2). These results indicate that higher shoot-to-root allocation is associated with stronger competitive effects in common wheat plants.

**Fig. 1.**
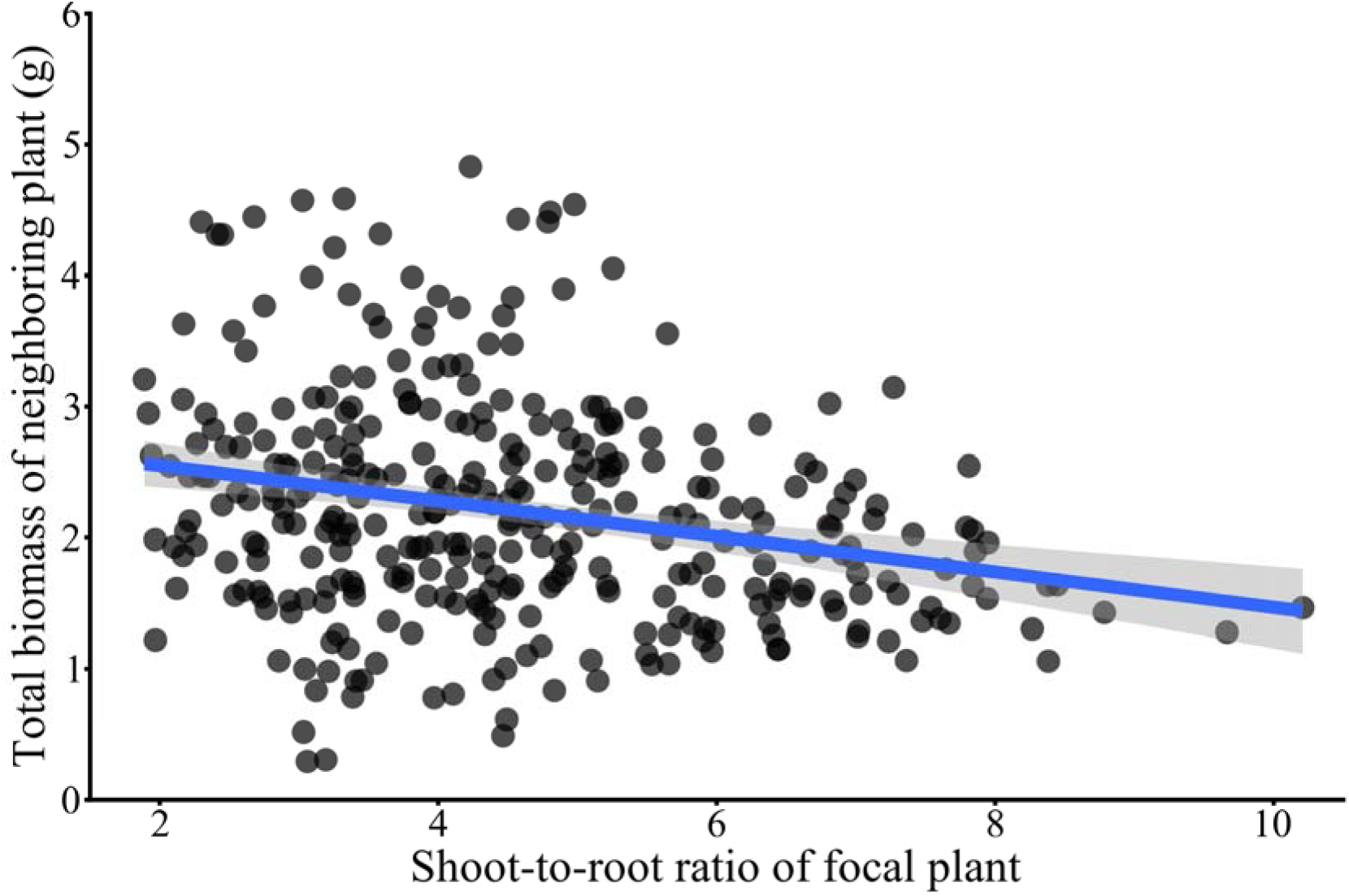
Relationship between the shoot-to-root ratio of focal plants and the total biomass of neighboring plants at 38 days. The line indicates significant correlations (LMM, *P* < 0.001). ALT TEXT: This figure showed effects of shoot-to-root ratio of focal wheat plants on total biomass of neighboring plants.

### Exp.2: Root-exudation application

The effects of root exudates on plant growth differed among cultivars (treatment × cultivar: d.f. = 11, *F* = 4.23, *P* < 0.001; Fig. S3). NBB and NP produced greater total biomass when exposed to root exudates from the same cultivar than when exposed to distilled water or exudates from different cultivars (Fig. S3), suggesting positive growth responses to cues from genetically similar neighbors. In contrast, the total biomass of N61 and CS did not differ among treatments (Fig. S3).

Cultivars also differed in their biomass allocation responses to root exudates (Fig. 2). The two landraces, NP and CS, showed no significant changes in shoot-to-root allocation across treatments (Fig. 2). In contrast, N61, the modern cultivar, exhibited little or no change in shoot-to-root allocation when exposed to root exudates from the same cultivar relative to distilled water (root × treatment, DW vs. Same: *F* = 0.025, *P* = 0.876), whereas shoot-to-root allocation tended to increase when exposed to root exudates from different cultivars (DW vs. Different: *F* = 4.927, *P* = 0.067; Same vs. Different: *F* = 4.156, *P* = 0.067; Fig. 2). These results suggest that N61 did not increase competitive allocation in response to same-cultivar exudates, while showing a tendency to increase allocation in response to different-cultivar exudates.

**Fig. 2.**
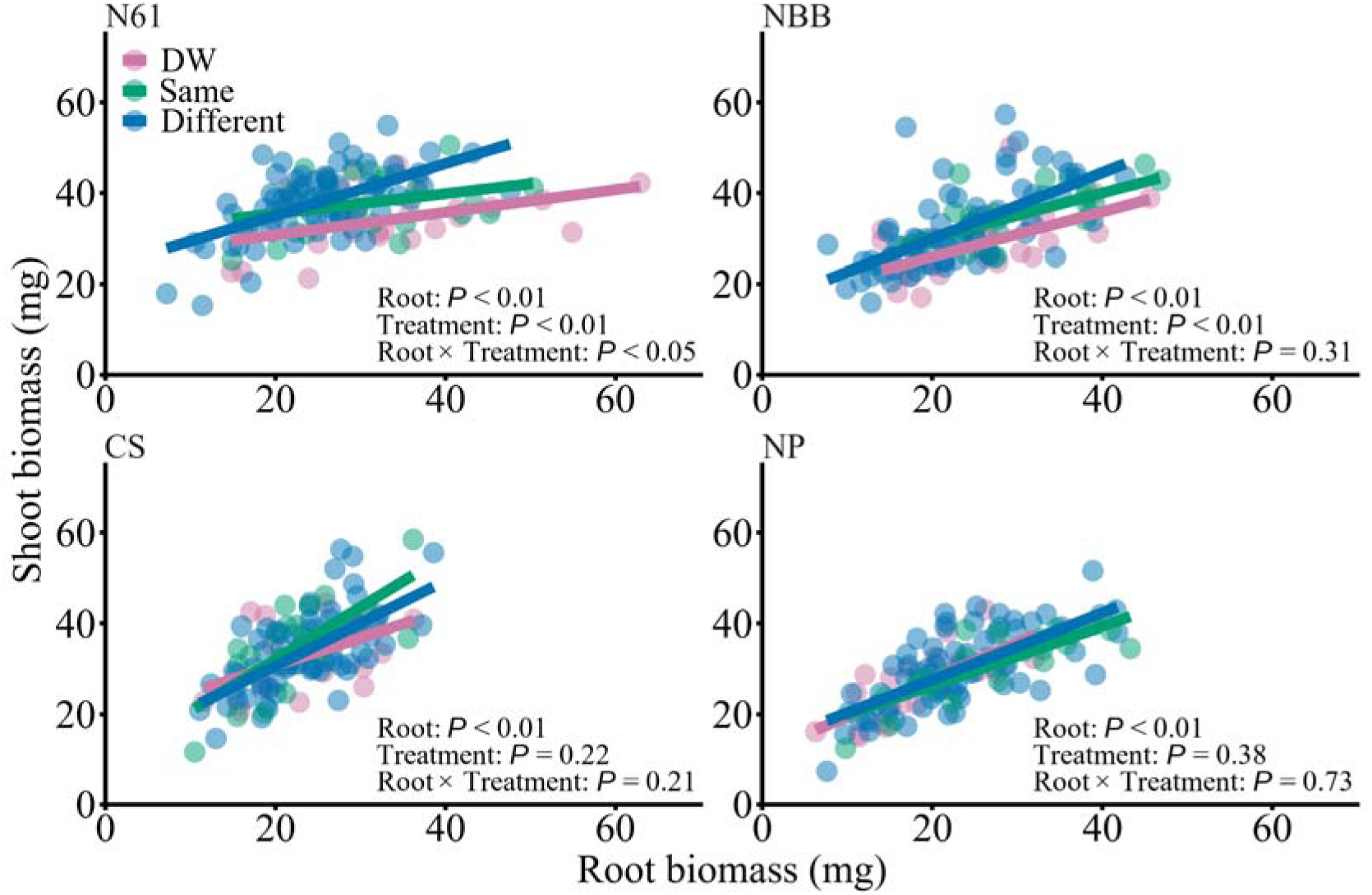
Effects of root biomass, treatment, and their interaction on shoot biomass of common wheat *Triticum aestivum* L. in the root exudate application experiment. Each cultivar seedling was exposed to distilled water (DW), the root exudates from the same cultivar (Same) or different cultivars (Different). N61, NBB, CS and NP stand for “Norin 61”, “Nobeokabouzu”, “Chinese Spring” and “KU-7113”, respectively. (GLM, *P* < 0.05) ALT TEXT: Figures showed shoot-to-root ratio of each cutlivar depending on treatments including distilled water and the root exudates from the same or different cultivars.

NBB showed a different pattern from N61. Shoot-to-root allocation increased in response to both same- and different-cultivar root exudates relative to distilled water (DW vs. Same: *F* = 7.02, *P* = 0.017; DW vs. Different: *F* = 10.30, *P* < 0.01), whereas no difference was detected between same- and different-cultivar exudates (Same vs. Different: *F* = 1.08, *P* = 0.302; Fig. 2). Thus, NBB appeared to respond to the presence of root-derived neighbor cues rather than to donor genotype.

Distilled-water treatments provided a baseline estimate of biomass allocation in the absence of neighbor-derived cues. Under this condition, baseline shoot-to-root allocation differed among cultivars, with NP and CS tending to show higher allocation than NBB and N61 (Fig. 3; Table S4). Moreover, N61 showed a gradual increase in shoot-to-root allocation as the genetic relatedness of neighboring cultivars decreased, resulting in allocation levels that varied according to neighbor identity (Fig. 4; Table S5). The same donor-identity-dependent pattern was not detected in the landraces (Fig. 2).

**Fig. 3.**
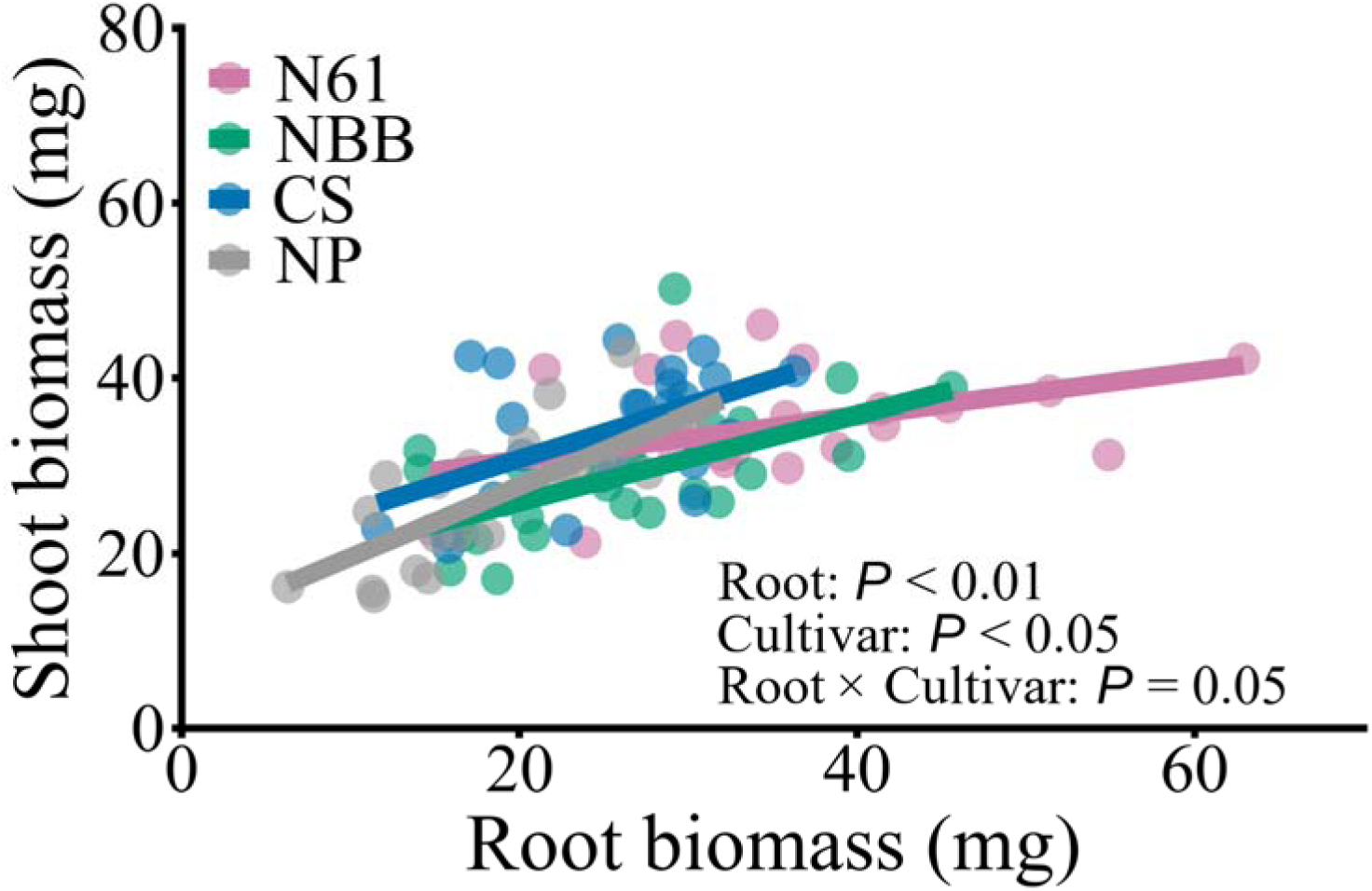
Effects of root biomass, cultivar, and their interaction on shoot biomass of common wheat *Triticum aestivum* L. cultivar in the root exudate application experiment. This shows only the results of distilled water treatment. N61, NBB, CS and NP stand for “Norin 61”, “Nobeokabouzu”, “Chinese Spring” and “KU-7113”, respectively. This analysis was used to evaluate baseline allocation patterns in the absence of neighbor-derived cues. ALT TEXT: This figure showed shoot-to-root ratio of each cultivar when exposed to distilled water. Norin 61, Nobeokabouzu, Chinese Spring and KU-7113 represent N61, NBB, CS and NP, respectively.

**Fig. 4.**
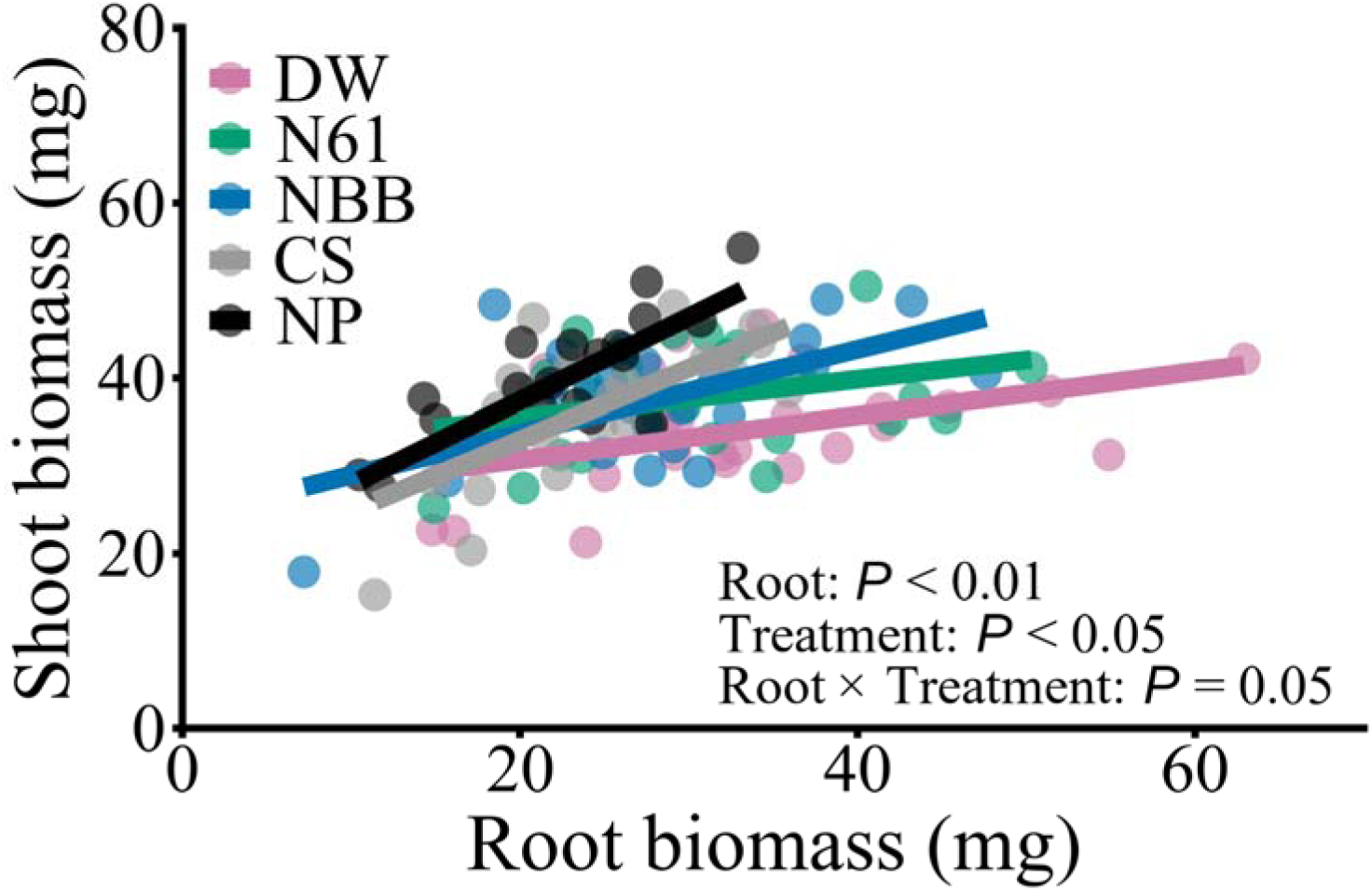
Relationships between root biomass and shoot biomass in Norin 61 (N61) receiver plants exposed to distilled water (DW) or root exudates from N61, NBB, CS, and NP in the root-exudate application experiment. Donor cultivars are ordered according to their genetic relatedness to N61 based on Takenaka et al. (2018). Differences in the slopes or intercepts of the relationships indicate treatment-dependent changes in shoot-to-root allocation. N61, NBB, CS, and NP stand for Norin 61, Nobeokabouzu, Chinese Spring, and KU-7113, respectively. ALT TEXT: This figure showed shoot-to-root ratio of Norin 61 depending on treatments including distilled water and the root exudates from the same or different cultivars.

### Exp.3: Yield measurements

Cultivars differed markedly in their several yield responses to competition (Figs. 5–6, Table S6). This pattern was also observed when heterospecific neighbors, *T. turgidum* L. subsp. *durum* (Desf.) Husnot cv. ‘Langdon’ was excluded from the analysis (Table S7). The modern cultivar Norin 61 (N61), which displayed genotype-dependent changes in shoot-to-root allocation in the root-exudate experiment, maintained grain biomass under same-cultivar competition but showed substantial yield reductions under different-cultivar competition. The three landraces showed cultivar-specific yield responses that did not consistently correspond to the genotype-dependent allocation pattern observed in N61.

**Fig. 5.**
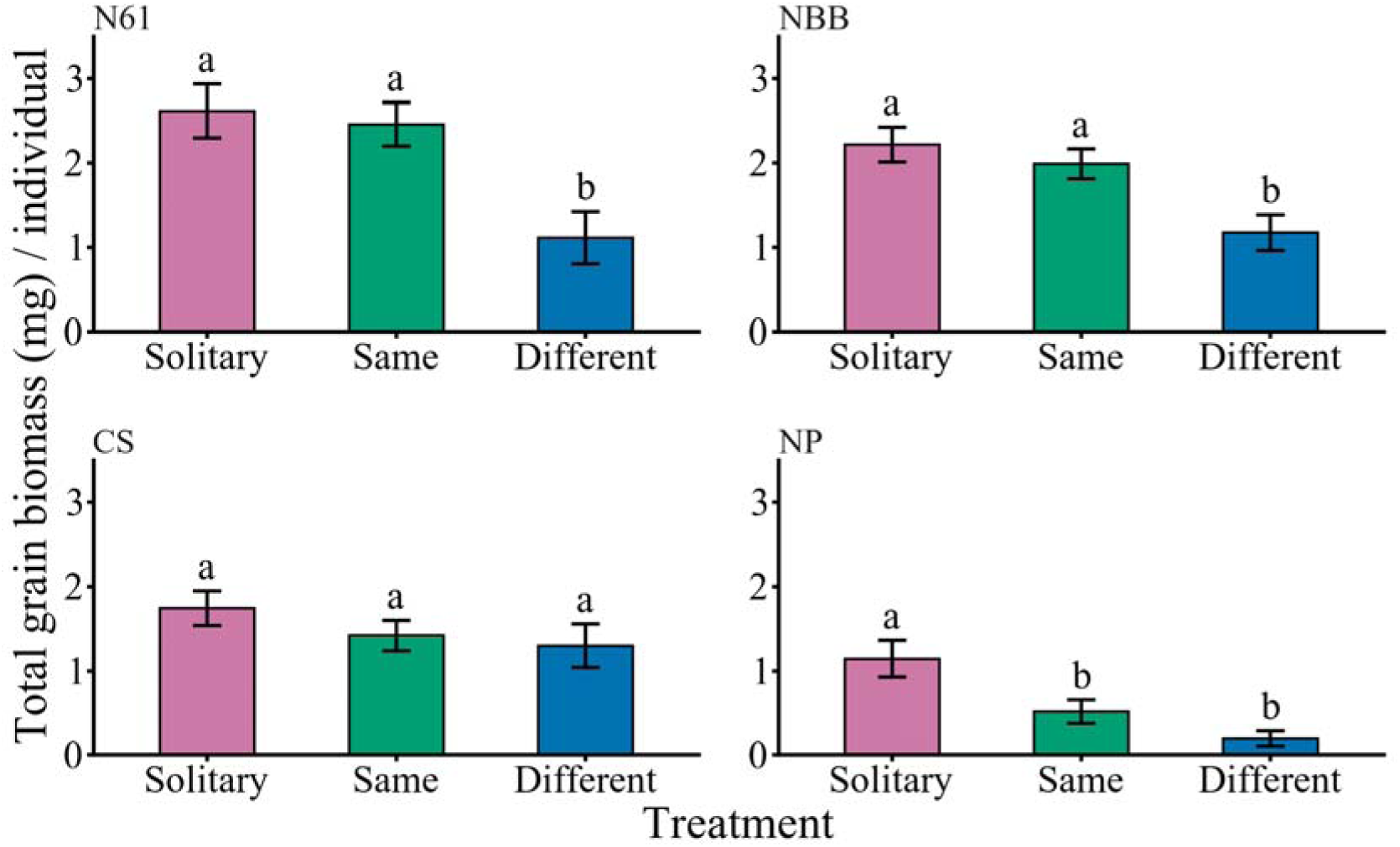
Total grain biomass per individual of each common wheat *Triticum aestivum* L. cultivar in each experimental condition: solitary (Solitary), the same cultivar (Same) and different cultivars (Different). N61, NBB, CS and NP stand for “Norin 61”, “Nobeokabouzu”, “Chinese Spring” and “KU-7113”, respectively. Bars represent SE. Different letters indicate significant differences among treatments within each cultivar (GLM, *P* < 0.05). ALT TEXT: Graphs showed total grain biomass of each cultivar depending on treatments including solitary, the same cultivar competition and different cultivars competition.

**Fig. 6.**
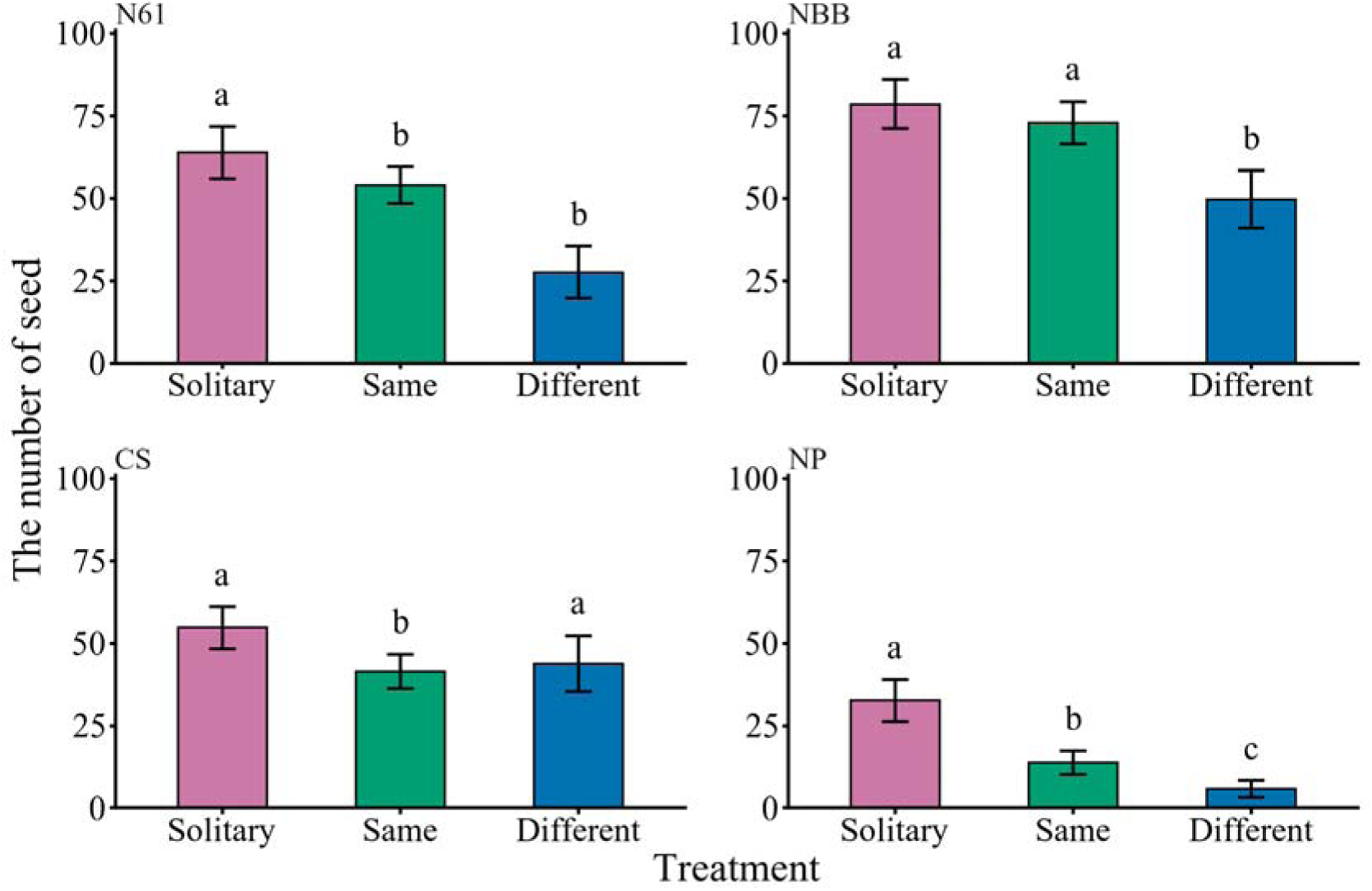
The number of grains of each common wheat *Triticum aestivum* L. cultivar in each experimental condition: solitary (Solitary), the same cultivar (Same) and different cultivars (Different). N61, NBB, CS and NP stand for “Norin 61”, “Nobeokabouzu”, “Chinese Spring” and “KU-7113”, respectively. Bars represent SE. Different letters indicate significant differences among treatments within each cultivar (GLM, *P* < 0.05). ALT TEXT: Graphs showed the number of grains of each cultivar depending on treatments including solitary, the same cultivar competition and different cultivars competition.

We measured major yield-related traits, including grain number, total grain biomass, panicles per plant, grains per panicle, thousand-grain weight, harvest index, and estimated field yield. Among these traits, total grain biomass showed the clearest responses to competition (Figs. 5–6). In N61, total grain biomass did not differ between solitary plants and plants grown with individuals of the same cultivar (Solitary vs. Same, χ^2^ = 0.097, *P* = 0.756; Fig. 5). However, total grain biomass was significantly reduced when plants competed with different cultivars (Solitary vs. Different, χ^2^ = 5.116, *P* = 0.036; Same vs. Different, χ^2^ = 5.878, *P* =0.036; Fig. 5). Grain number declined in the presence of competitors (Solitary vs. Same, χ^2^= 24.399, *P* < 0.001; Solitary vs. Different, χ^2^ = 7.904, *P* = 0.007; Same vs. Different, χ^2^= 0.412, *P* = 0.521; Fig. 6). On average, N61 retained approximately 94% of the total grain biomass observed in solitary plants when grown with individuals of the same cultivar, whereas grain biomass declined to approximately 43% of solitary levels under different-cultivar competition. Harvest index (HI), which is the ratio of grain yield to biological yield or biomass (Donald and Hamblin, 1976; Hay, 1995), also differed among treatments in N61, with lower values under different-cultivar competition than under solitary or same-cultivar conditions (Solitary vs. Same, *F* = 0.006, *P* = 0.937; Solitary vs. Different, *F* = 5.40, *P* = 0.035; Same vs. Different, *F* = 7.51, *P* = 0.022; Table S8). In contrast, HI did not differ among neighboring treatments in NBB and CS (NBB, *F* = 0.014, *P* = 0.986; CS, *F* = 0.107, *P* = 0.898), but lower HI in NP in the presence of neighbors (NP, Solitary vs. Same, *F* = 4.95, *P* = 0.043; Solitary vs. Different, *F* = 5.37, *P* = 0.043; Same vs. Different, *F* = 0.303, *P* = 0.583; Table S8).

The landraces showed different patterns from N61. In NP, both total grain biomass and grain number were reduced in the presence of competitors (grain biomass: Solitary vs. Same, χ^2^ = 4.035, *P* = 0.067; Solitary vs. Different, χ^2^ = 5.413, *P* = 0.060; grain number: Solitary vs. Same, χ^2^ = 4.651, *P* = 0.031; Solitary vs. Different, χ^2^ = 78.318, *P* < 0.001; Fig. 5, 6). Total grain biomass was reduced to approximately 45% of solitary levels under same-cultivar competition and to approximately 17% under different-cultivar competition. CS exhibited reduced grain numbers under same-cultivar competition (Solitary vs. Same, χ^2^ = 7.294, *P* = 0.010; Same vs. Different, ^2^ = 12.612, *P* = 0.003; Fig. 6), although total grain biomass did not differ significantly among treatments (χ^2^ = 1.059, *P* = 0.589; Fig. 5).

NBB also maintained total grain biomass under same-cultivar competition but showed reduced total grain biomass under different-cultivar competition. Total grain biomass was significantly lower in different-cultivar treatments than in solitary or same-cultivar treatments (Solitary vs. Same, χ^2^ = 0.453, *P* = 0.501; Solitary vs. Different, χ^2^ = 8.088, *P* = 0.013; Same vs. Different, χ^2^ = 5.436, *P* = 0.030; Fig. 5). Grain number also declined in different-cultivar competition than in solitary and the same-cultivar competition (Solitary vs. Same, χ^2^ = 0.877, *P* = 0.349; Solitary vs. Different, χ^2^ = 20.027, *P* < 0.001; Same vs. Different, χ^2^ = 16.472, *P* < 0.001; Fig. 6). Thus, NBB showed a yield-maintenance pattern under same-cultivar competition that was partly similar to N61, despite showing no clear genotype-dependent allocation response in the root-exudate experiment. Thus, N61 showed the clearest correspondence between genotype-dependent allocation responses and yield maintenance under same-cultivar competition. However, the yield response of NBB indicates that yield maintenance under same-cultivar competition may also arise through mechanisms other than genotype-dependent reduction of shoot-to-root allocation.

## Discussion

Our findings suggest that genotype-dependent responses may contribute to yield maintenance in common wheat plants, particularly in the modern cultivar Norin 61 (N61), and raise the possibility that modern breeding may have favored conditional regulation of competition rather than uniformly reduced competitiveness. The modern wheat cultivar examined in this study, N61, exhibited reduced competitive allocation toward genetically similar neighbors while tending to maintain competitive responses toward genetically distinct cultivars. Consistent with this pattern, N61 maintained grain biomass more effectively under same-cultivar competition than under competition with different cultivars. However, the response of Nobeokabouzu (NBB) indicates that yield maintenance under same-cultivar competition is not necessarily explained by genotype-dependent reduction of shoot-to-root allocation in all cultivars. In contrast, the other landraces showed either no detectable genotype-dependent allocation responses or yield responses that were not clearly associated with genotype-dependent allocation. Together, these results suggest that genotype discrimination can contribute to crop productivity by reducing self-competition among genetically similar individuals in monoculture stands, while also indicating that common wheat cultivars may use multiple mechanisms to regulate competition and maintain yield.

Our first competition experiment demonstrated that shoot-to-root allocation functions as a competitive trait in common wheat plants. Individuals with higher shoot-to-root allocation suppressed the growth of neighboring plants more strongly, regardless of whether competitors belonged to the same or different cultivars or species. Previous studies have frequently used changes in shoot-to-root allocation as a common indicator of kin or genotype discrimination resulting from kin or genotype recognition because this trait reflects investment in competitive resource acquisition (Dudley and File, 2007; Dudley *et al*., 2013; Yang *et al*., 2018; Fukano *et al*., 2019). Our results provide direct evidence that variation in shoot-to-root allocation is associated with competitive effects on neighboring individuals in common wheat plants, validating its use as a proxy for competitive allocation in subsequent experiments.

Cultivars differed markedly in their responses to root exudates. The landraces NP and CS showed no detectable changes in shoot-to-root allocation across treatments, suggesting either an absence of genotype-dependent responses or responses that were weaker than those observed in N61. In contrast, NBB altered its allocation pattern when exposed to neighboring plants regardless of whether the root exudates originated from the same or different cultivars. This pattern suggests that NBB responds to the presence of neighboring plants but does not discriminate among neighboring cultivars. N61 exhibited a distinct response pattern: shoot-to-root allocation remained unchanged when exposed to root exudates from the same cultivar compared to distilled water but tended to increase when exposed to exudates from different cultivars. Moreover, allocation increased progressively as the genetic relatedness of neighboring cultivars decreased. These findings suggest that N61 adjusts competitive allocation according to neighbor identity rather than simply responding to the presence of neighboring plants.

### Breeding may favor conditional competitiveness rather than reduced competitiveness

Classical crop ideotype theory proposes that breeding improves productivity by reducing individual competitiveness (Donald, 1968; Weiner, 2003). Under this view, modern cultivars are expected to exhibit weaker competitive responses than their ancestral counterparts. However, our results suggest a more nuanced mechanism. N61 did not appear to lose competitive ability altogether. Instead, it suppressed competitive allocation when interacting with individuals of the same cultivar while tending to maintain competitive responses toward genetically distinct cultivars. This pattern is consistent with a strategy of conditional competitiveness, in which competitive traits are expressed selectively according to social context (Dudley *et al*. 2013: Yamawo 2026). Such a strategy could allow plants to avoid unnecessary allocation to competition within monoculture stands while retaining the capacity to compete when encountering genetically distinct neighbors. Although only a single modern cultivar was examined in this study, the observed pattern raises the possibility that modern breeding may favor conditional rather than uniformly reduced competitiveness.

The baseline allocation patterns observed under distilled-water treatment further support this interpretation, although they should be interpreted cautiously because not all pairwise cultivar differences were statistically significant (Table S4). NP and CS exhibited relatively high shoot-to-root allocation even in the absence of neighbor-derived cues, whereas N61 exhibited substantially lower allocation. Interestingly, the allocation level of N61 exposed to root exudates from genetically distant cultivars approached the baseline allocation levels observed in NP and CS. This pattern suggests that landraces may maintain a constitutively competitive phenotype, whereas N61 expresses competitive allocation more plastically according to social context. Such plasticity may reduce the costs associated with unnecessary competition among genetically similar neighbors.

The yield experiment provides support for the ecological consequences of this strategy in N61. The most striking result was that N61 retained approximately 94% of the grain biomass observed in solitary plants when grown with individuals of the same cultivar. In contrast, grain biomass declined to approximately 43% of solitary levels when N61 competed with different cultivars. This more than two-fold difference suggests that suppressing competitive allocation toward genetically identical neighbors may substantially reduce intraspecific competition with the same cultivar and contribute to yield maintenance under monoculture stands. Furthermore, N61 did not change harvest index, the value expressing biomass allocation from shoot to seed, in the presence of the same cultivar, but exhibit lower it in the presence of different cultivars. This indicates that N61 preferentially allocates the acquired resources to competitive traits, namely aboveground growth in the presence of genetically distinct neighbors. As the expression of competitive traits entails allocation costs, this investment likely reduced the resources available for reproduction, leading to lower seed production in different cultivar neighboring treatment. Therefore, N61 exhibited little reduction in several yield-related traits under same-cultivar competition, whereas substantially larger reductions occurred under different-cultivar competition.

In contrast, the landraces provided weaker evidence that genotype-dependent responses contributed to yield maintenance. NP exhibited strong reductions in grain biomass and grain number under both same- and different-cultivar competition, consistent with its lack of detectable genotype-dependent responses. NBB maintained grain biomass under same-cultivar competition, but its root-exudate responses did not clearly depend on donor genotype. This discrepancy suggests that yield maintenance under same-cultivar competition is not necessarily mediated by genotype-dependent reduction of shoot-to-root allocation in all cultivars. Instead, NBB may maintain yield through responses to neighbor presence, differences in intrinsic competitive tolerance, or compensatory allocation to reproductive traits. CS maintained relatively stable grain biomass across treatments despite reductions in grain number in the presence of the same cultivar, indicating that additional factors may influence competitive outcomes in this cultivar. Together, these results suggest that the capacity to reduce competition through genotype-dependent responses varies substantially among common wheat cultivars, and that genotype-dependent reduction represents one possible mechanism rather than a universal explanation for yield maintenance.

Common wheat cultivars appear to span a continuum from constitutive competitiveness to genotype-dependent competitiveness. At one end of this continuum, the landraces NP and CS appeared to show weak or undetectable genotype-dependent allocation responses, suggesting a constitutively competitive strategy. At the opposite end, the modern cultivar N61 adjusted competitive allocation according to the genetic identity of neighboring plants, suppressing competition toward individuals of the same cultivar while tending to maintain responses toward genetically distinct cultivars. NBB occupied an intermediate position, responding to the presence of neighboring plants but showing little evidence of discrimination among cultivars. These patterns suggest that variation among cultivars is not simply a matter of the presence or absence of genotype discrimination, but rather reflects different strategies for balancing competitive ability and resource-use efficiency. More broadly, this continuum suggests that artificial selection may modify not only the strength of competitiveness itself but also the degree to which competitive traits are deployed conditionally according to social context.

Plants can use multiple cues to discriminate among neighboring individuals, including volatile organic compounds, light profiles, and root-derived chemical cues (Biedrzycki *et al*., 2010; Karban *et al*., 2013; Semchenko *et al*., 2014; Crepy and Casal, 2015; Yamawo *et al*., 2017; Yang *et al*., 2018; Shiojiri *et al*., 2021; Chen *et al*., 2023; Ito *et al*., 2025). Our root-exudate experiment demonstrated that common wheat cultivars can alter competitive allocation in response to belowground cues alone, indicating that root exudates play an important role in genotype discrimination or neighbor-cue responses in common wheat plants. However, we did not investigate the potential contributions of aboveground cues. Future studies integrating volatiles, light-mediated, and root-derived signals will be necessary to obtain a more comprehensive understanding of neighbor discrimination mechanisms in common wheat.

Our findings also contribute to a growing recognition that kin or genotype discrimination varies substantially within species. Many studies implicitly assume that all individuals within a species exhibit similar discrimination abilities. However, recent work has highlighted the importance of intraspecific variation in recognition and discrimination mechanisms (Anten and Chen, 2021; Mazal *et al*., 2023). The strong differences observed among common wheat cultivars demonstrate that discrimination ability is not uniform within a species and may itself be an evolving trait. Such variation provides opportunities to identify the genetic and physiological mechanisms underlying plant discrimination systems through comparative and quantitative genetic approaches.

Several authors have suggested that kin or genotype discrimination should decline under artificial selection because crop breeding favors reduced competitiveness and greater uniformity (Kiers and Denison, 2014; Fukano *et al*., 2019). However, empirical studies, including the present work, repeatedly report genotype-dependent responses in modern crop cultivars (Ninkovic, 2003; Fang *et al*., 2013; Zhu and Zhang, 2013; Murphy *et al*., 2017*b*; Yang *et al*., 2018; Saleh and Kniss, 2022). One possible explanation is that genotype discrimination remains advantageous because crops interact not only with conspecific neighbors but also with weeds and other heterospecific competitors. Under such conditions, the ability to suppress competition among genetically identical neighbors while retaining competitive responses toward unrelated individuals may increase productivity. In fact, only one study demonstrated that several rice cultivars inhibit the growth of heterospecific competitors, weeds, in the paddy field when planted with genetically similar neighbors (Xu *et al*., 2021). Although there are few examples of this hypothesis being tested, it offers an intriguing explanation for the persistence of genotype-dependent responses despite strong artificial selection.

### Limitations and future directions

First, only a single modern cultivar, Norin 61 (N61), was included in the experiments. Therefore, our results demonstrate that genotype-dependent reduction of competition can occur in a modern wheat cultivar, but they do not establish that this is a general feature of modern cultivars. Moreover, cultivar category and genetic background were confounded in the present design. The response observed in N61 may therefore reflect N61-specific alleles or genetic background rather than the effect of modern breeding per se. Cultivars carrying the same causal allele, or functionally equivalent alleles, may exhibit similar responses irrespective of whether they are classified as landraces or modern cultivars. Testing the effects of breeding history will require comparisons among multiple independently derived modern cultivars and appropriate ancestral or pedigree-matched landraces, together with genetic mapping or association analyses. Second, genotype recognition was inferred from cultivar-specific responses, genotype discrimination, to root exudates and associated changes in biomass allocation rather than from direct measurements of recognition mechanisms. Future studies combining physiological, metabolomic, and genetic approaches will be required to identify the molecular basis of genotype recognition and its phenotypic manifestation as genotype discrimination in common wheat plants. Third, our experiments were conducted under controlled conditions using pairwise interactions among cultivars. Because agricultural fields contain more complex competitive environments, including variation in planting density, resource availability, and weed pressure, further studies under field conditions will be needed to evaluate the ecological importance of genotype recognition and discrimination in crop production systems. Fourth, because NBB maintained grain biomass under same-cultivar competition without showing clear genotype-dependent allocation responses, future studies should examine alternative mechanisms of yield maintenance, such as competitive tolerance, reproductive compensation, and responses to general neighbor-derived cues. Despite these limitations, the consistent association between genotype-dependent allocation patterns and yield maintenance in N61 suggests that conditional suppression of competition among genetically similar neighbors may represent a previously underappreciated mechanism contributing to productivity in monoculture crops.

In conclusion, we demonstrated that (1) shoot-to-root allocation functions as a competitive trait in common wheat plants; (2) the modern cultivar N61 suppresses competitive allocation toward individuals of the same cultivar while tending to maintain responses toward genetically distinct cultivars; and (3) this genotype-dependent adjustment of competitive allocation is associated with substantially greater yield maintenance under monoculture conditions in N61. These findings suggest that genotype discrimination potentially mediated by genotype recognition may contribute to crop productivity by reducing intraspecific competition among genetically identical individuals. At the same time, the response of NBB indicates that yield maintenance can also arise through cultivar-specific mechanisms other than genotype-dependent suppression of shoot-to-root allocation. More broadly, our results raise the possibility that modern crop breeding may favor conditional competitiveness, allowing modern crop species to balance cooperation with genetically similar neighbors and competition with genetically distinct individuals. Because only a single modern cultivar was examined in this study, future work incorporating a broader range of modern cultivars will be necessary to determine how widespread this strategy is among contemporary common wheat cultivars. Our results suggest that conditional competitiveness may represent a previously overlooked target of crop breeding.

## Acknowledgements

The Authors thank the National BioResource Project (NBRP) Wheat for providing the common wheat cultivar seeds. This work was supported by the Joint Research Program of the Center for Ecological Research, Kyoto University, Japan.

## Author contributions

Nao Kosugi: Conceptualization, Formal analysis, Investigation, Writing-Original Draft Preparation. Takuto Kaneko: Formal analysis, Validation, Funding Acquisition, Visualization, Writing-Review & Editing. Kentaro Yoshida: Resource, Writing-Review & Editing. Akira Yamawo: Conceptualization, Formal analysis, Investigation, Validation, Funding Acquisition, Visualization, Project Administration, Supervision, Writing-Original Draft Preparation.

## Data availability

All datasets are available from the corresponding authors upon reasonable request.

## Supplements

**Table S1.**
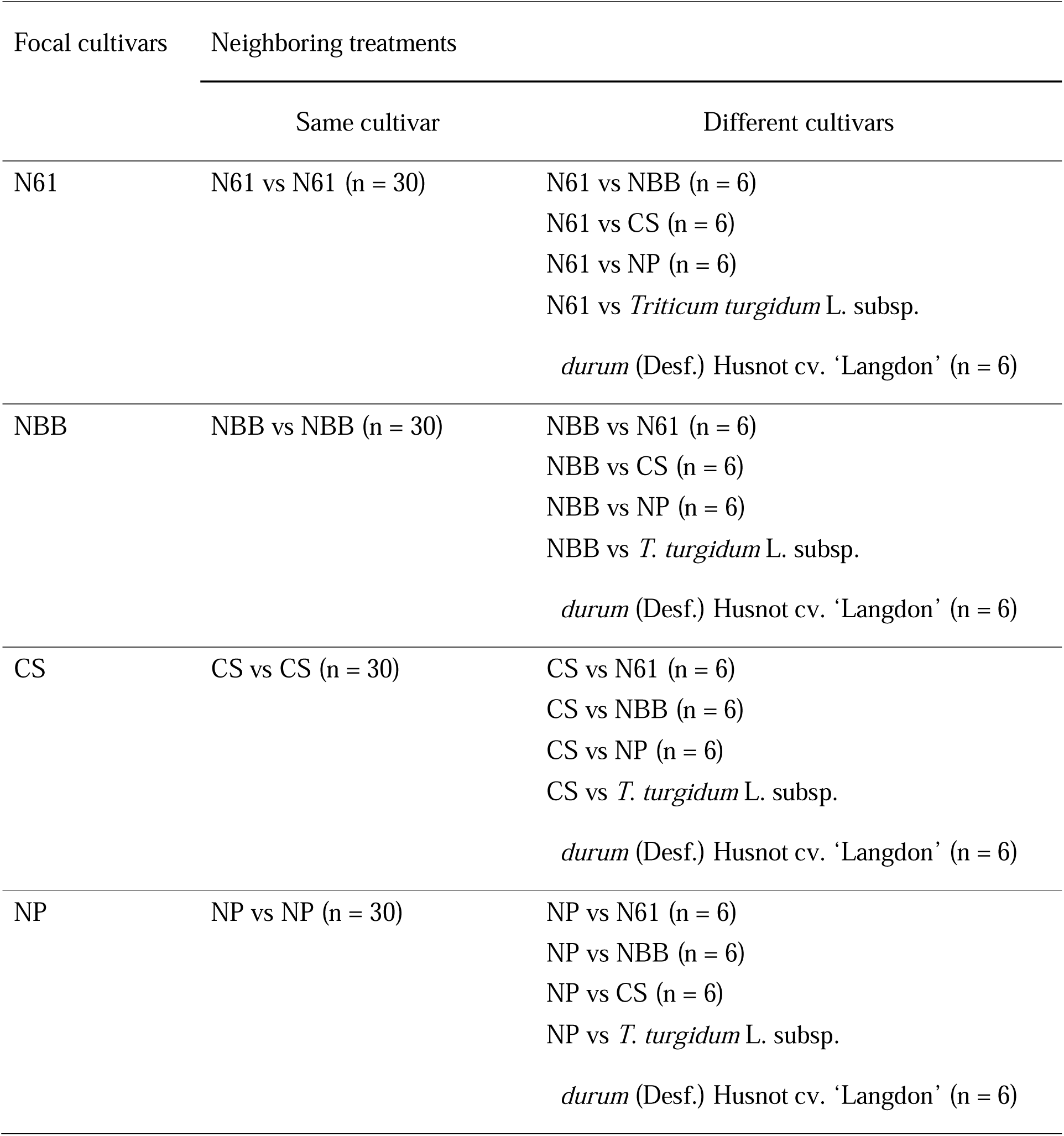
Pairs of first competition experiment.

**Table S2.**
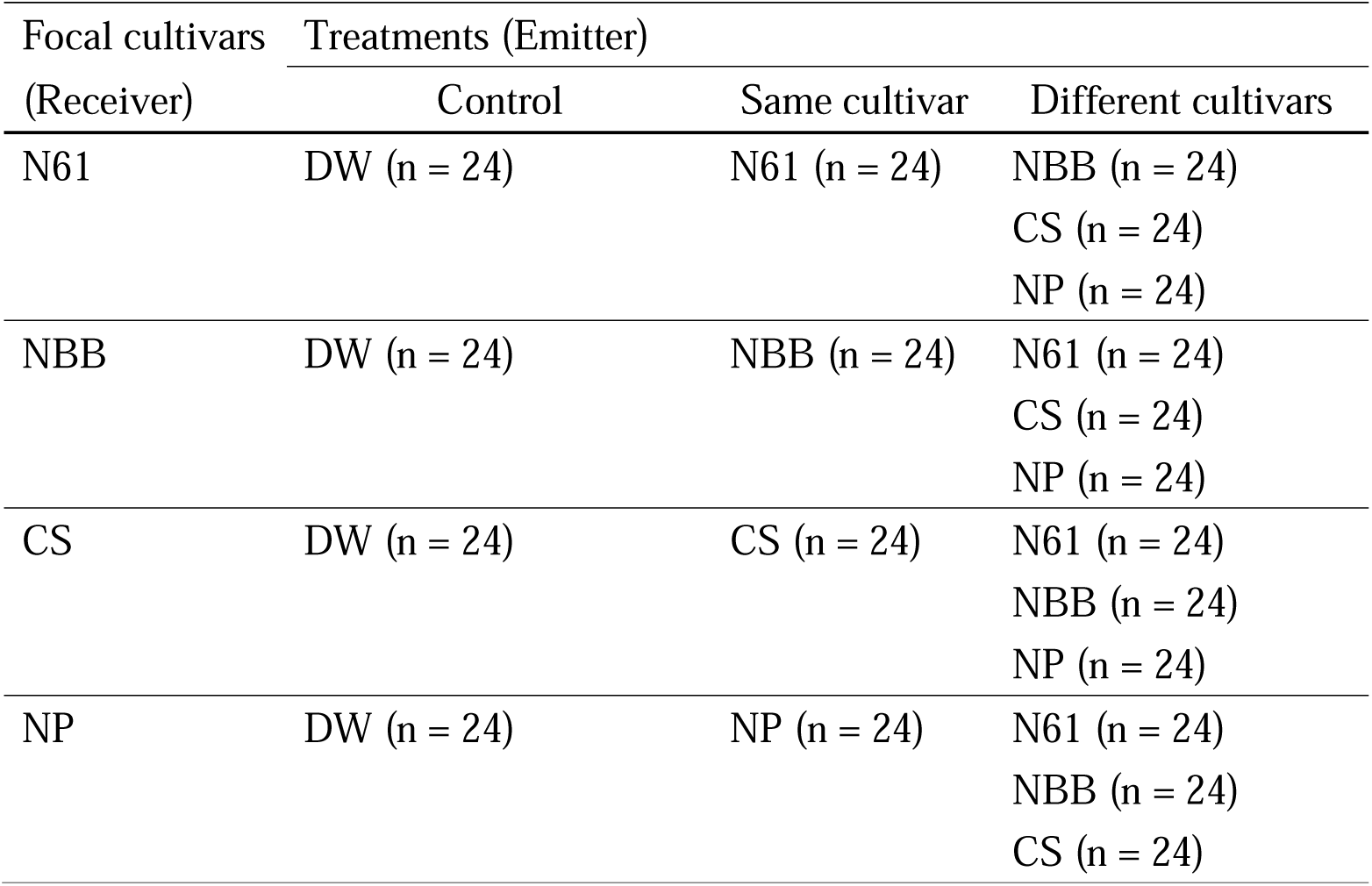
Emitter and receiver plants of the root exudate application experiment.

**Table S3.**
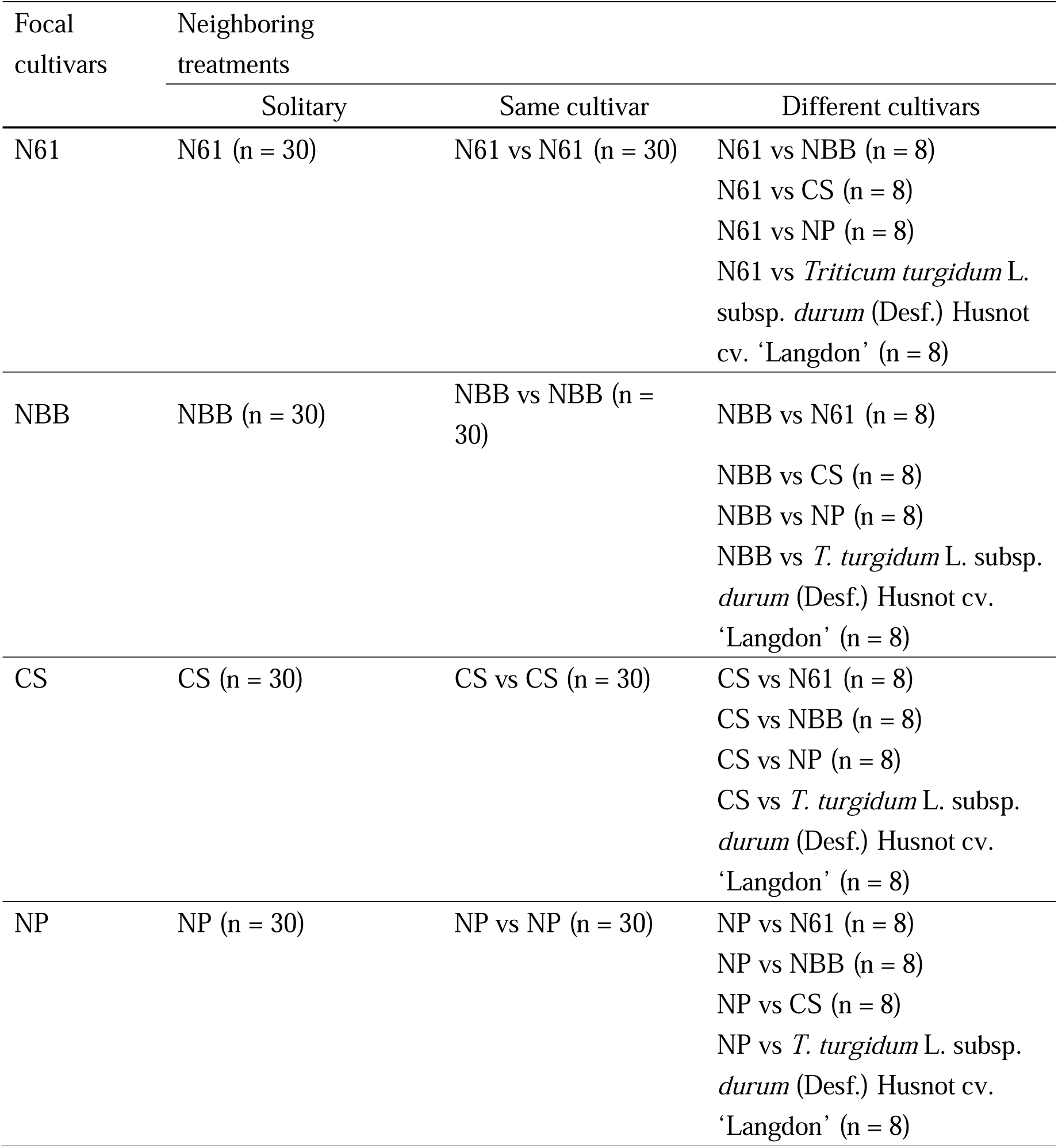
Pairs of second competition experiment.

**Table S4.**
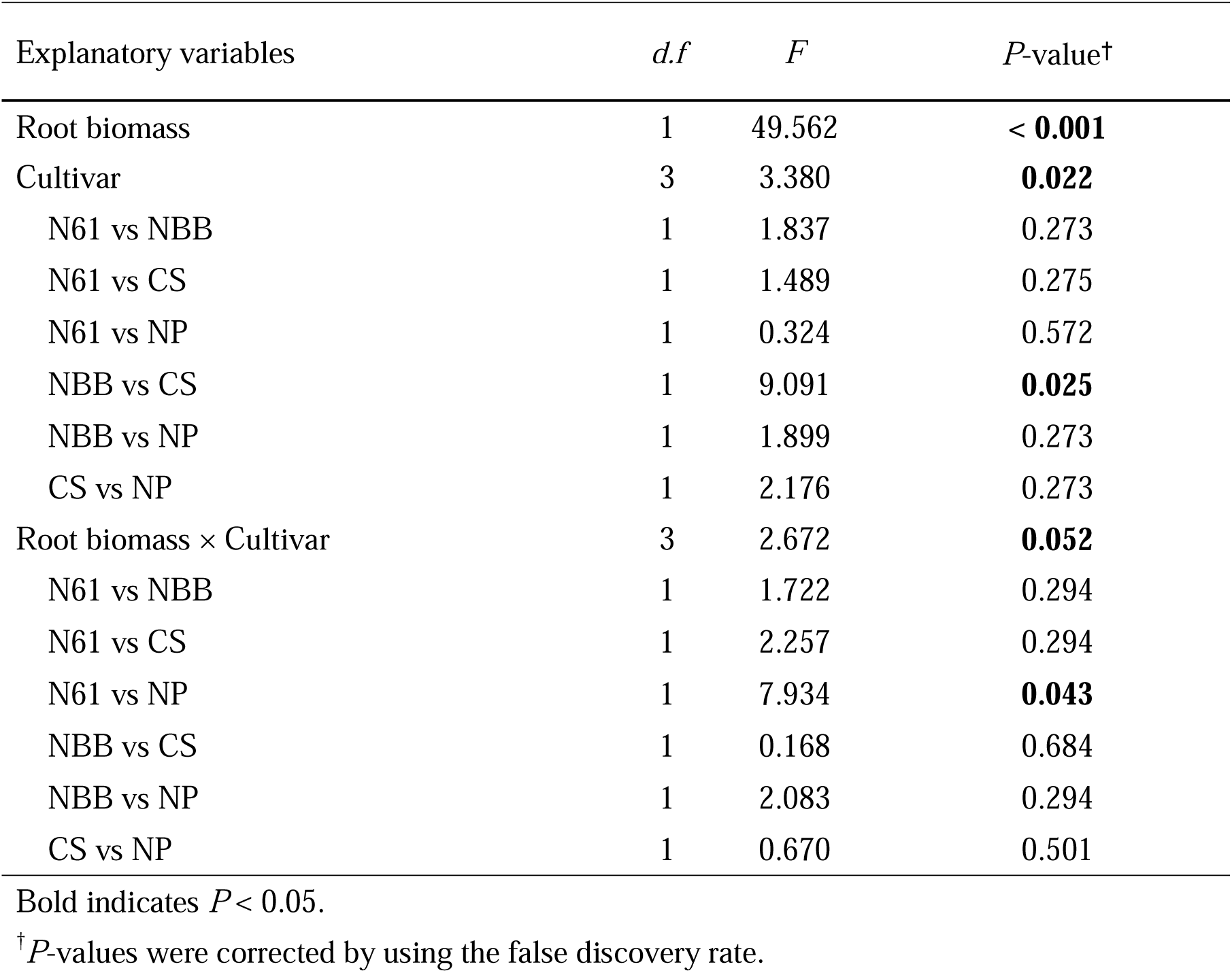
Results of generalized linear model examining the effect of root biomass, cultivar and their interaction on shoot biomass in distilled water treatment.

**Table S5.**
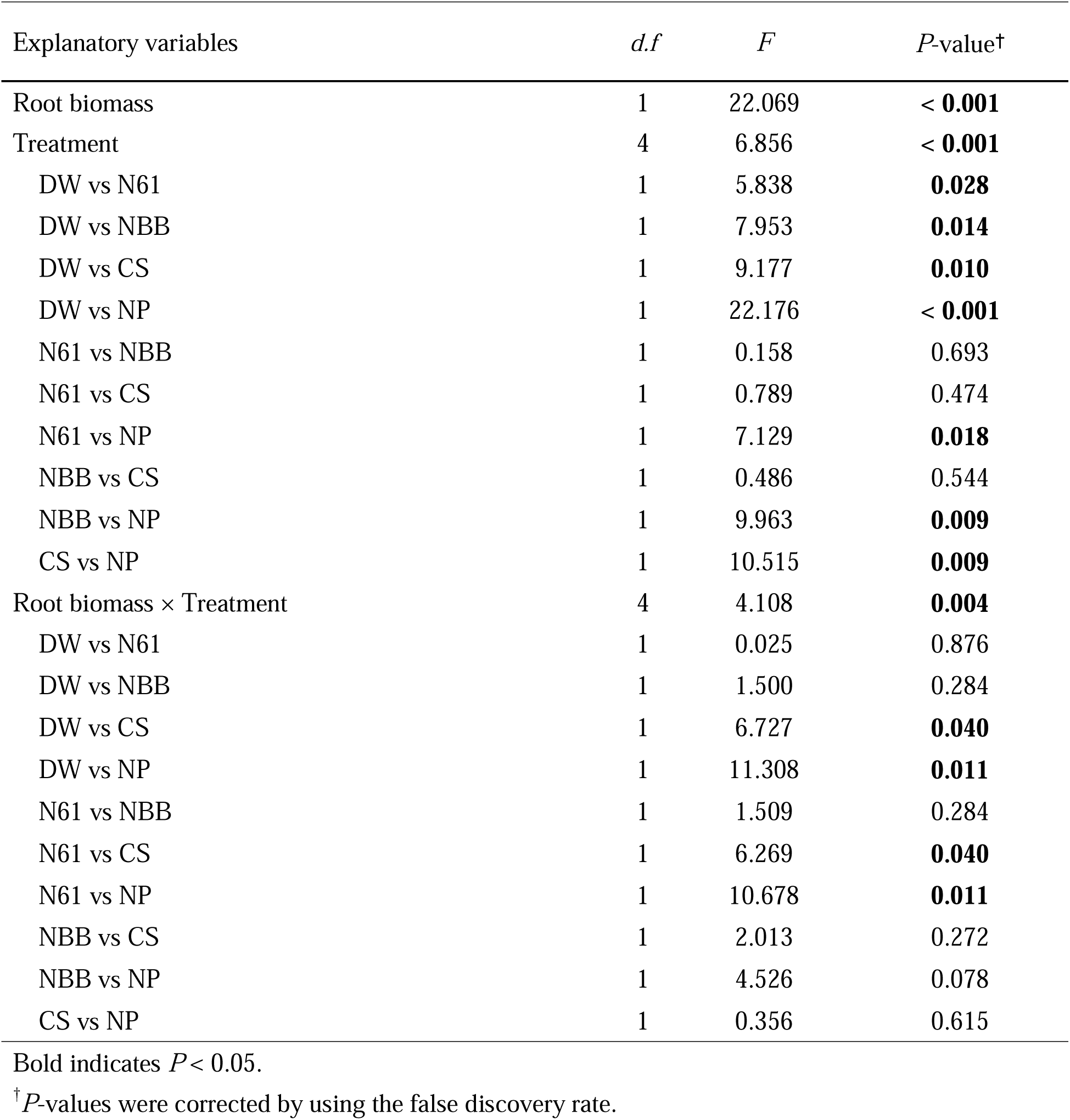
Results of generalized linear model examining the effect of root biomass, treatment and their interaction on shoot biomass in Norin 61.

**Table S6.**
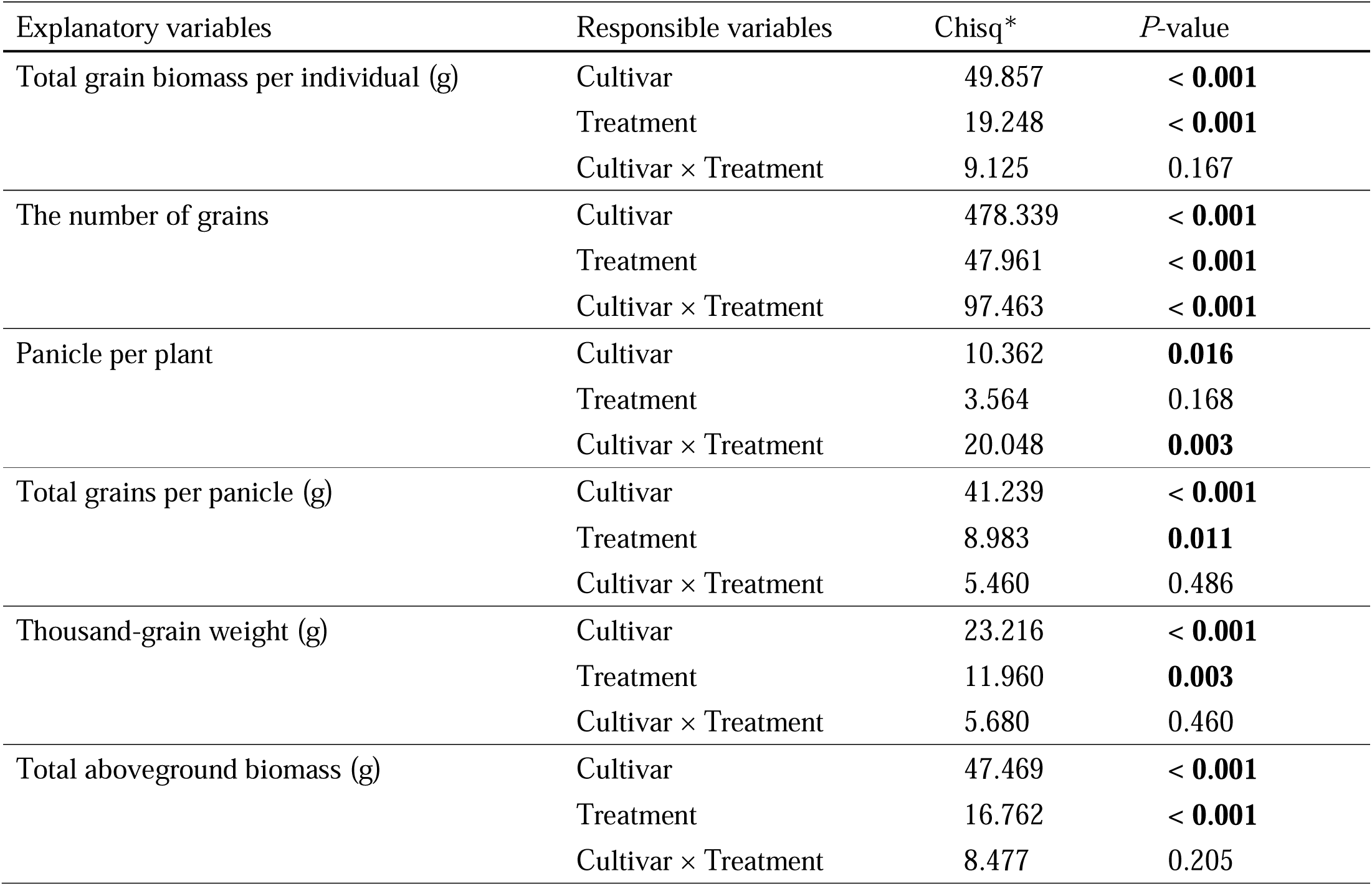
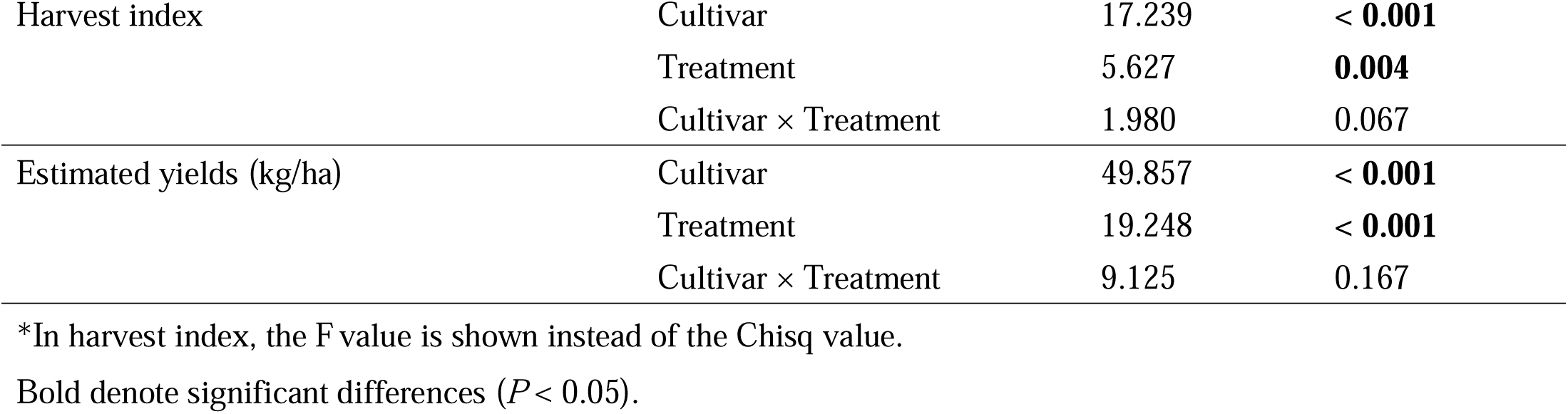
Results of analysis by generalized liner model, with each yield parameter as the responsible variables, cultivar, treatment and their interaction as the explanatory variables.

**Table S7.**
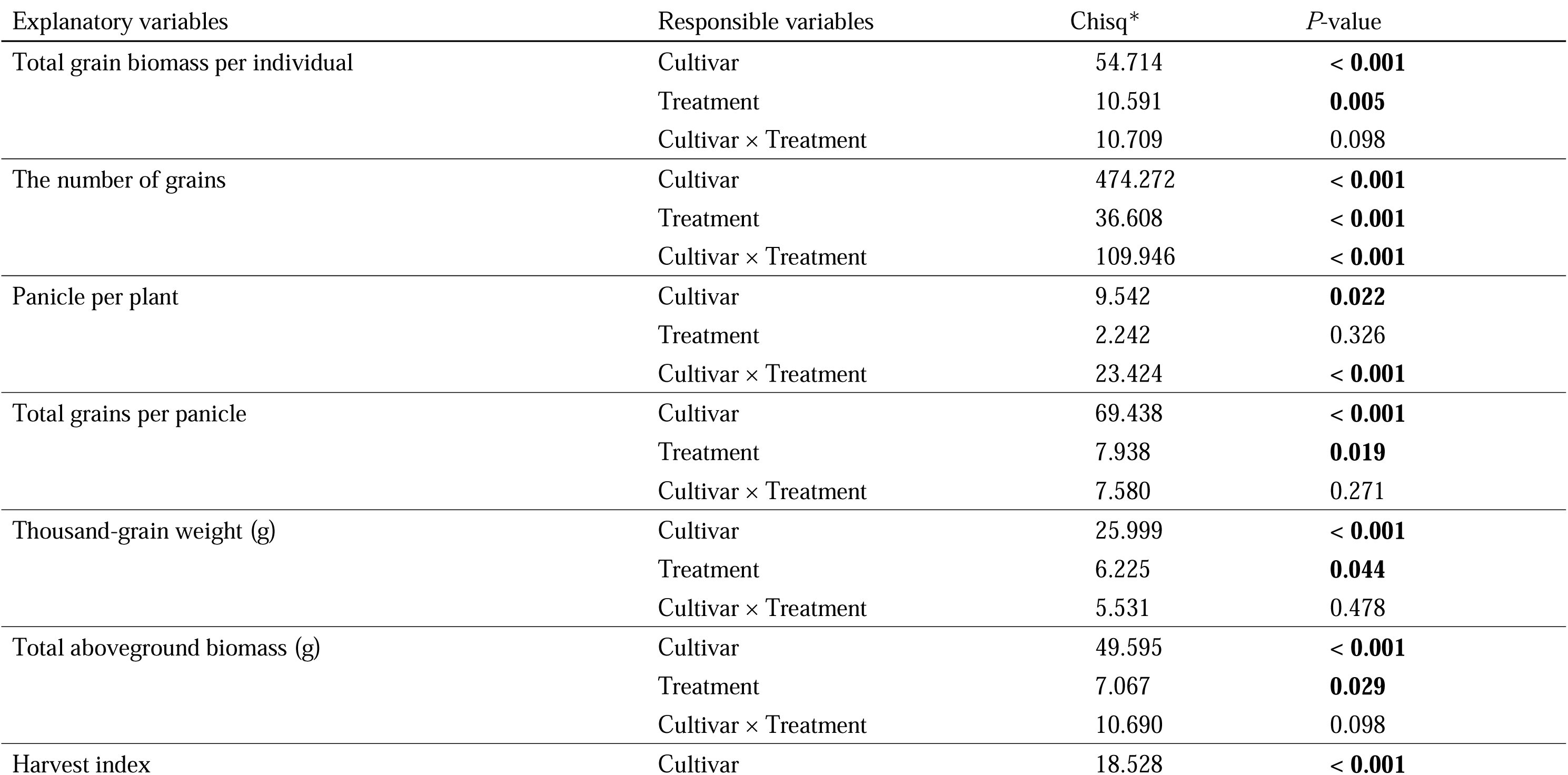
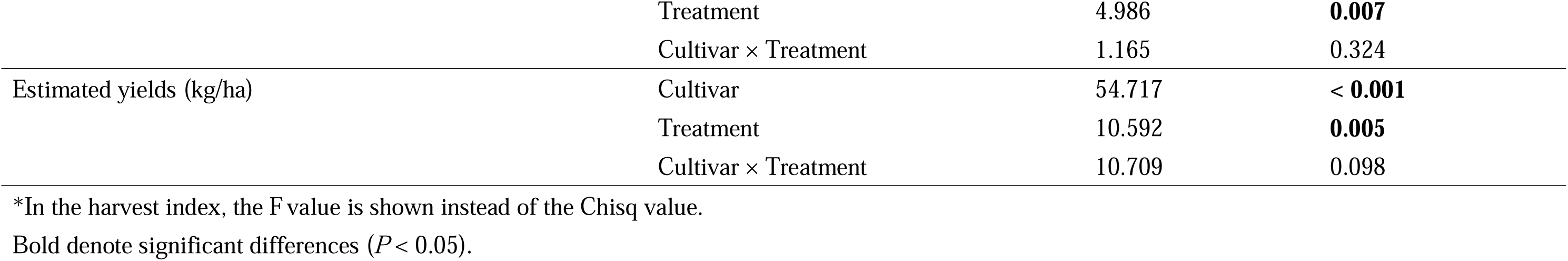
Results except for heterospecific, *Triticum turgidum* L. subsp. *durum* (Desf.) Husnot cv. ‘Langdon’ of analysis by generalized liner model, with each yield parameter as the responsible variables, cultivar, treatment and their interaction as the explanatory variables.

**Table S8.**
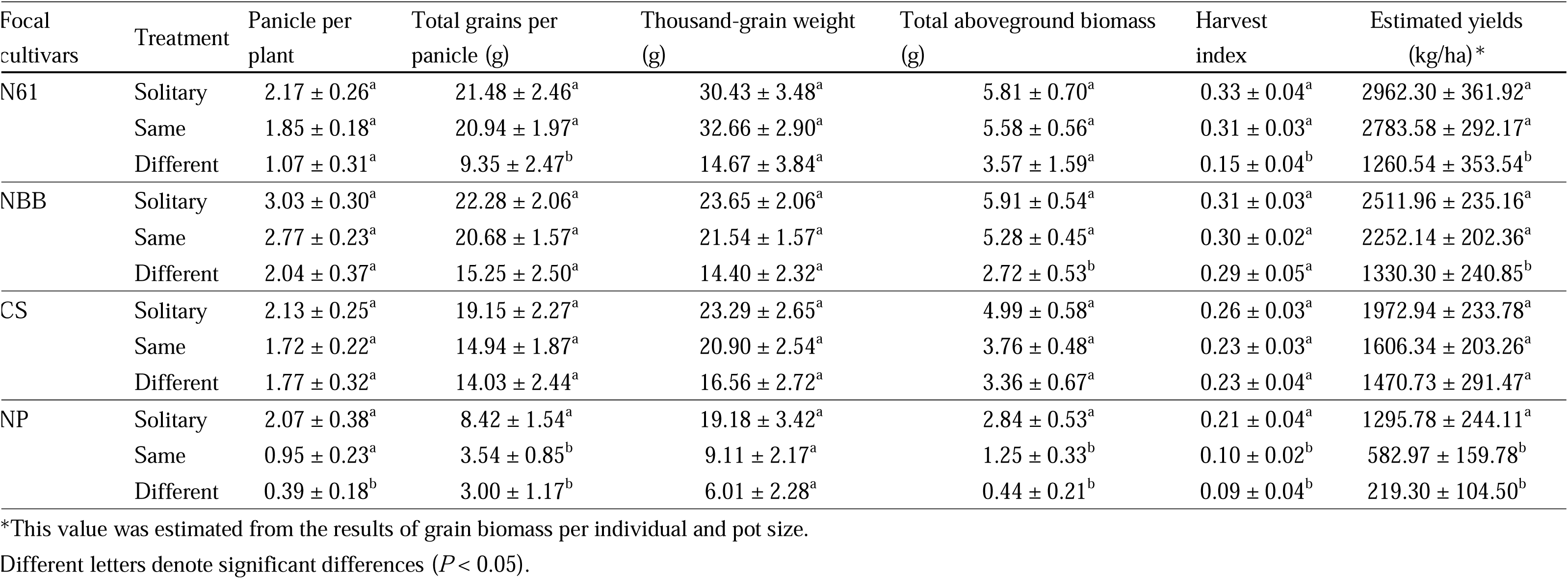
Yield parameters of each focal cultivar across neighboring treatments.

**Fig. S1.**
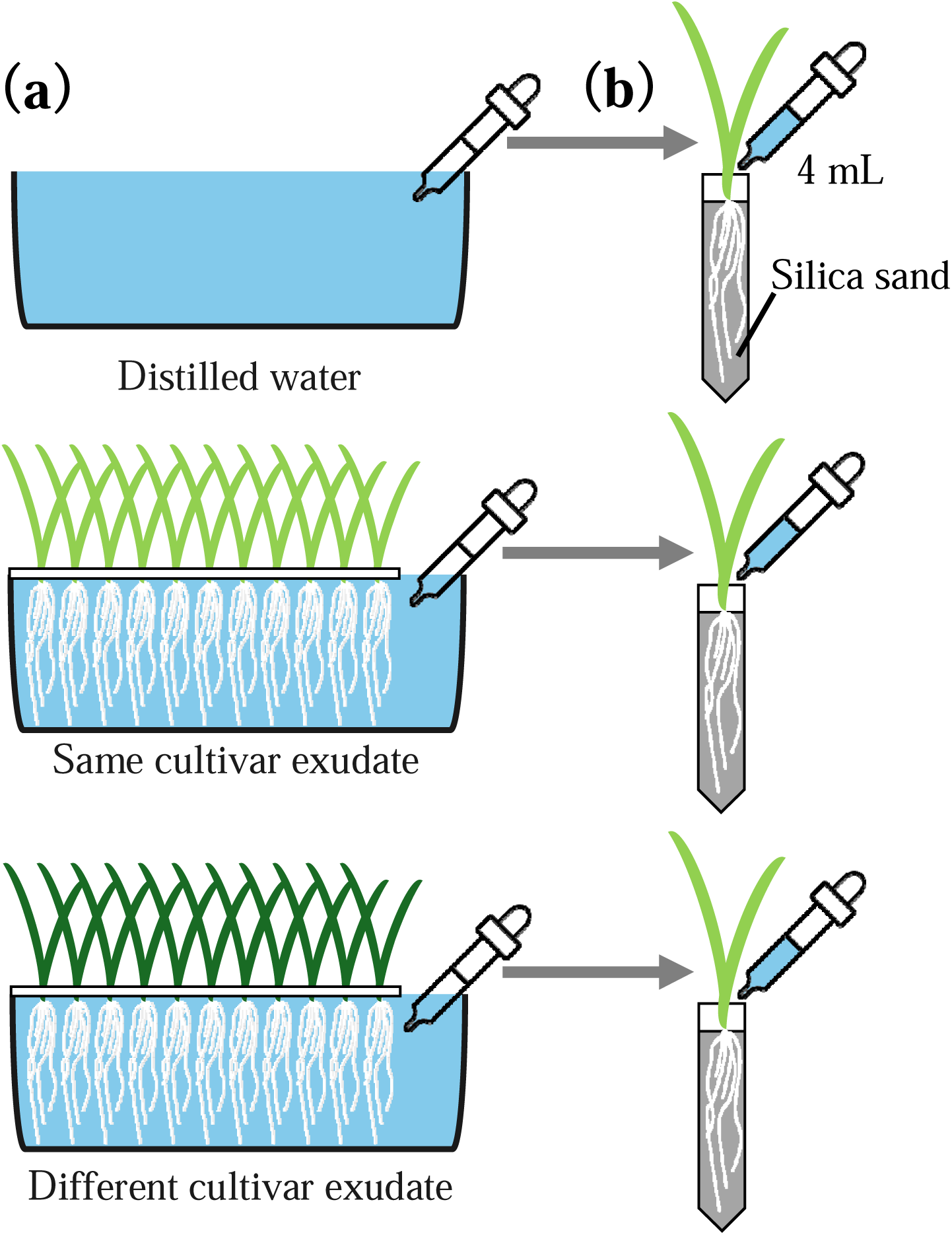
Methods of root exudate collection and application. **(a)** Root exudate collection: fifty seedlings of each common wheat *Triticum aestivum* L. cultivar were grown under hydroponic conditions with 300mL distilled water. **(b)** Root exudate application: 4 mL distilled water or the root exudates of the same cultivar or different cultivars were applied to seedlings of each cultivar transplanted into a 10 mL pipette tip containing silica sand every day. ALT TEXT: Figures showed (a) root exudate collection and (b) application 4mL of root exudate.

**Fig. S2.**
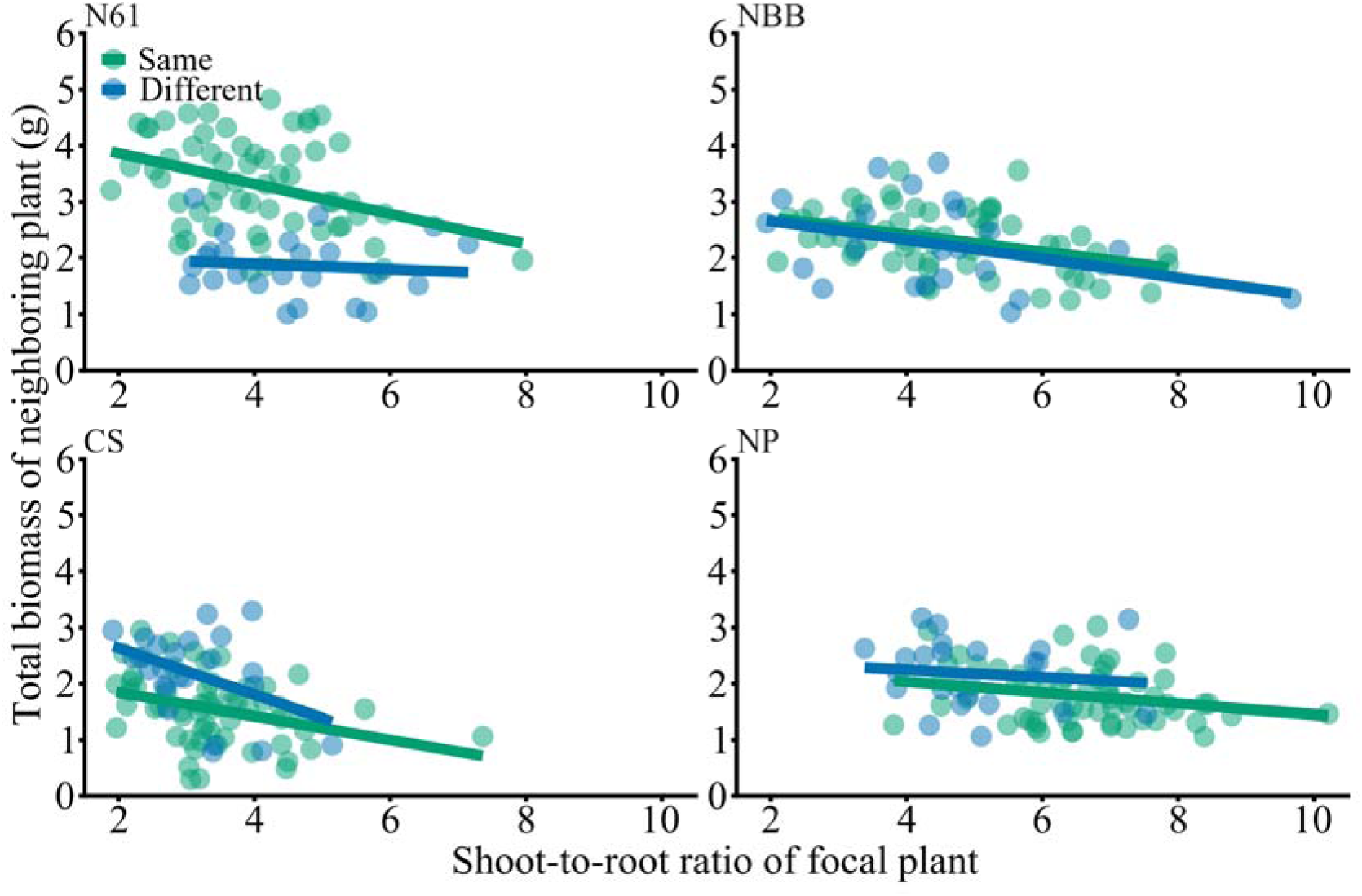
Relationship between shoot-to-root allocation and competitor biomass of each common wheat cultivar at 38 days. Each line indicates significant correlations (LMM, *P*< 0.001). ALT TEXT: This figure showed effects of shoot-to-root ratio of focal wheat plants on total biomass of neighboring plants for neighboring treatments.

**Fig. S3.**
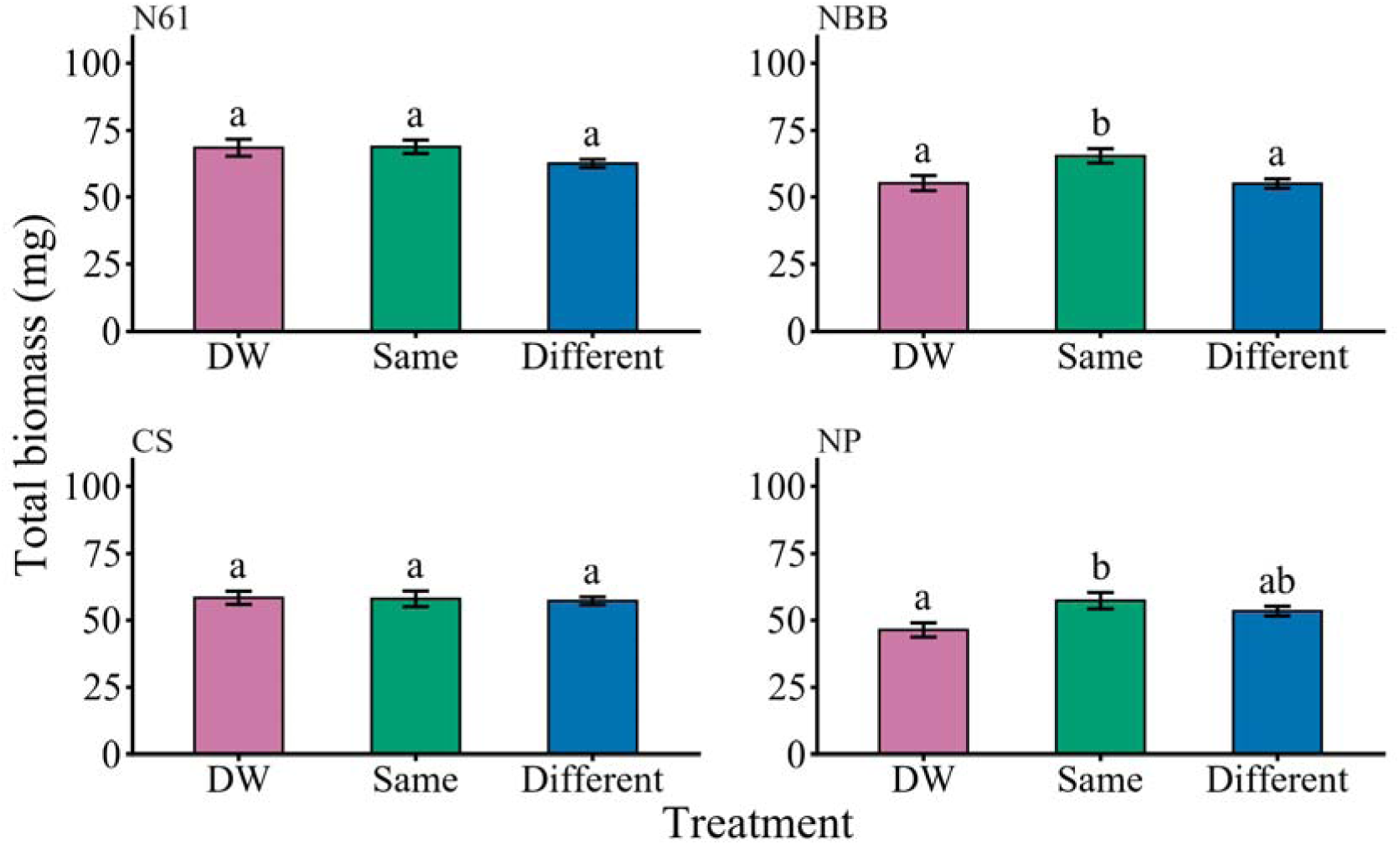
Total biomass of each common wheat *Triticum aestivum L.* cultivar in each experimental condition: exposed to distilled water (DW), the root exudates from the same cultivar (Same) or different cultivars (Different). N61, NBB, CS and NP stand for “Norin 61”, “Nobeokabouzu”, “Chinese Spring” and “KU-7113”, respectively. Bars represent SE. Different letters show significant differences (GLM, *P* < 0.05). ALT TEXT: Graphs showed total biomass of each cultivar depending on root exudate treatments.

## References

Anten NPR, Chen BJW. 2021. Detect thy family: Mechanisms, ecology and agricultural aspects of kin recognition in plants. Plant, Cell & Environment 44, 1059–1071.

Biedrzycki ML, Jilany TA, Dudley SA, Bais HP. 2010. Root exudates mediate kin recognition in plants. Communicative & Integrative Biology 3, 28–35.

Chen BJW, During HJ, Anten NPR. 2012. Detect thy neighbor: Identity recognition at the root level in plants. Plant Science 195, 157–167.

Chen L, Pan Y, Pan X, Yu L, Sosa A, Yang J, Li B. 2023. From competition to cooperation: Kin selection against selfish shade avoidance behaviour promotes plant invasions. Journal of Ecology 111, 645–654.

Crepy MA, Casal JJ. 2015. Photoreceptor mediated kin recognition in plants. New Phytologist 205, 329–338.

Donald CM. 1968. The breeding of crop ideotypes. Euphytica 17, 385–403.

Donald CM, Hamblin J. 1976. The Biological Yield and Harvest Index of Cereals as Agronomic and Plant Breeding Criteria. Advances in Agronomy. Elsevier, 361–405.

Dudley SA, File AL. 2007. Kin recognition in an annual plant. Biology Letters 3, 435–438.

Dudley SA, Murphy GP, File AL. 2013. Kin recognition and competition in plants. (D Robinson, Ed.). Functional Ecology 27, 898–906.

Fang S, Clark RT, Zheng Y, Iyer-Pascuzzi AS, Weitz JS, Kochian LV, Edelsbrunner H, Liao H, Benfey PN. 2013. Genotypic recognition and spatial responses by rice roots. Proceedings of the National Academy of Sciences 110, 2670–2675.

Fang S, Gao X, Deng Y, Chen X, Liao H. 2011. Crop Root Behavior Coordinates Phosphorus Status and Neighbors: From Field Studies to Three-Dimensional in Situ Reconstruction of Root System Architecture. Plant Physiology 155, 1277–1285.

Fukano Y, Guo W, Noshita K, Hashida S, Kamikawa S. 2019. Genotype aggregated planting improves yield in Jerusalem artichoke (*Helianthus tuberosus*) due to self/non self discrimination. Evolutionary Applications 12, 508–518.

Gersani M, Brown JS, O’Brien EE, Maina GM, Abramsky Z. 2001. Tragedy of the commons as a result of root competition. Journal of Ecology 89, 660–669.

Hay RKM. 1995. Harvest index: a review of its use in plant breeding and crop physiology. Annals of Applied Biology 126, 197–216.

Ito K, Ohsaki H, Novoplansky A, Hirota SK, Yamawo A. 2025. Integrated above- and below-ground interplant cueing of salt stress. Plant Signaling & Behavior 20, 2542560.

Karban R, Shiojiri K, Ishizaki S, Wetzel WC, Evans RY. 2013. Kin recognition affects plant communication and defence. Proceedings of the Royal Society B: Biological Sciences 280, 20123062.

Kiers ET, Denison RF. 2014. Inclusive fitness in agriculture. Philosophical Transactions of the Royal Society B: Biological Sciences 369, 20130367.

Maina GG, Brown JS, Gersani M. 2002. Intra-plant versus Inter-plant Root Competition in Beans: avoidance, resource matching or tragedy of the commons. Plant Ecology 160, 235–247.

Mazal L, Fajardo A, Till Bottraud I, Corenblit D, Fumanal B. 2023. Kin selection, kin recognition and kin discrimination in plants revisited: A claim for considering environmental and genetic variability. Plant, Cell & Environment 46, 2007–2016.

Murphy GP, Dudley SA. 2007. Above and below ground competition cues elicit independent responses. Journal of Ecology 95, 261–272.

Murphy GP, Swanton CJ, Van Acker RC, Dudley SA. 2017*a*. Kin recognition, multilevel selection and altruism in crop sustainability. (D Gibson, Ed.). Journal of Ecology 105, 930–934.

Murphy GP, Van Acker R, Rajcan I, Swanton CJ. 2017*b*. Identity recognition in response to different levels of genetic relatedness in commercial soya bean. Royal Society Open Science 4, 160879.

Ninkovic V. 2003. Volatile communication between barley plants affects biomass allocation. Journal of Experimental Botany 54, 1931–1939.

Novoplansky A. 2019. What plant roots know? Seminars in Cell & Developmental Biology 92, 126–133.

O’Brien EE, Gersani M, Brown JS. 2005. Root proliferation and seed yield in response to spatial heterogeneity of below ground competition. New Phytologist 168, 401–412.

Saleh OS, Kniss AR. 2022. The growth behaviour of winter wheat (*Triticum aestivum* L.) in the presence of inter- and intraspecific neighbours. Canadian Journal of Plant Science 102, 1053–1056.

Sedgley RH. 1991. An appraisal of the Donald ideotype after 21 years. Field Crops Research 26, 93–112.

Semchenko M, Saar S, Lepik A. 2014. Plant root exudates mediate neighbour recognition and trigger complex behavioural changes. New Phytologist 204, 631–637.

Sher J, Zheng Y, Burns JH, Jan G, Zhang J-L. 2025. Kin recognition in plants-an ecological perspective: an overview of plant kin recognition under different resources, consequences and future challenges. Journal of Plant Interactions 20, 2548579.

Shiojiri K, Ishizaki S, Ando Y. 2021. Plant–plant communication and community of herbivores on tall goldenrod. Ecology and Evolution 11, 7439–7447.

Takenaka S, Nitta M, Nasuda S. 2018. Population structure and association analyses of the core collection of hexaploid accessions conserved *ex situ* in the Japanese gene bank NBRP-Wheat. Genes & Genetic Systems 93, 237–254.

Takigahira H, Yamawo A. 2019. Competitive responses based on kin-discrimination underlie variations in leaf functional traits in Japanese beech (Fagus crenata) seedlings. Evolutionary Ecology 33, 521–531.

Weiner J. 2003. Ecology – the science of agriculture in the 21st century. The Journal of Agricultural Science 141, 371–377.

Xu Y, Cheng H, Kong C, Meiners SJ. 2021. Intra specific kin recognition contributes to inter specific allelopathy: A case study of allelopathic rice interference with paddy weeds. Plant, Cell & Environment 44, 3709–3721.

Yamawo A. 2015. Relatedness of Neighboring Plants Alters the Expression of Indirect Defense Traits in an Extrafloral Nectary-Bearing Plant. Evolutionary Biology 42, 12–19.

Yamawo A. 2021. Intraspecific competition favors ant–plant protective mutualism. Plant Species Biology 36, 372–378.

Yamawo A. 2026. Kin discrimination in plants: overview and implications for population and community ecology. Biological Reviews doi: 10.1002/brv.70194.

Yamawo A, Mukai H. 2020. Outcome of interspecific competition depends on genotype of conspecific neighbours. Oecologia 193, 415–423.

Yamawo A, Sato M, Mukai H. 2017. Experimental evidence for benefit of self discrimination in roots of a clonal plant. AoB PLANTS 9.

Yang X-F, Li L-L, Xu Y, Kong C-H. 2018. Kin recognition in rice (Oryza sativa) lines. New Phytologist 220, 567–578.

Zhu L, Zhang D-Y. 2013.Donald’s Ideotype and Growth Redundancy: A Pot Experimental Test Using an Old and a Modern Spring Wheat Cultivar. (I De Smet, Ed.). PLoS ONE 8, e70006.

