## Supplementary Files for "Genotype-dependent reductions of competition are associated with yield maintenance in a modern common wheat *Triticum aestivum* L. cultivar"

**Supplements**

| Table S1. Pairs of first competition experiment | |  |
| --- | --- | --- |
| Focal cultivars | Neighboring treatments |  |
|  | Same cultivar | Different cultivars |
| N61 | N61 vs N61 (n = 30) | N61 vs NBB (n = 6) |
|  |  | N61 vs CS (n = 6) |
|  |  | N61 vs NP (n = 6) |
|  |  | N61 vs *Triticum turgidum* L. subsp. |
|  |  | *durum* (Desf.) Husnot cv. ‘Langdon’ (n = 6) |
| NBB | NBB vs NBB (n = 30) | NBB vs N61 (n = 6) |
|  |  | NBB vs CS (n = 6) |
|  |  | NBB vs NP (n = 6) |
|  |  | NBB vs *T*. *turgidum* L. subsp. |
|  |  | *durum* (Desf.) Husnot cv. ‘Langdon’ (n = 6) |
| CS | CS vs CS (n = 30) | CS vs N61 (n = 6) |
|  |  | CS vs NBB (n = 6) |
|  |  | CS vs NP (n = 6) |
|  |  | CS vs *T*. *turgidum* L. subsp. |
|  |  | *durum* (Desf.) Husnot cv. ‘Langdon’ (n = 6) |
| NP | NP vs NP (n = 30) | NP vs N61 (n = 6) |
|  |  | NP vs NBB (n = 6) |
|  |  | NP vs CS (n = 6) |
|  |  | NP vs *T*. *turgidum* L. subsp. |
|  |  | *durum* (Desf.) Husnot cv. ‘Langdon’ (n = 6) |

| Table S2. Emitter and receiver plants of the root exudate application experiment | | | |
| --- | --- | --- | --- |
| Focal cultivars | Treatments (Emitter) |  |  |
| (Receiver) | Control | Same cultivar | Different cultivars |
| N61 | DW (n = 24) | N61 (n = 24) | NBB (n = 24) |
|  |  |  | CS (n = 24) |
|  |  |  | NP (n = 24) |
| NBB | DW (n = 24) | NBB (n = 24) | N61 (n = 24) |
|  |  |  | CS (n = 24) |
|  |  |  | NP (n = 24) |
| CS | DW (n = 24) | CS (n = 24) | N61 (n = 24) |
|  |  |  | NBB (n = 24) |
|  |  |  | NP (n = 24) |
| NP | DW (n = 24) | NP (n = 24) | N61 (n = 24) |
|  |  |  | NBB (n = 24) |
|  |  |  | CS (n = 24) |

| Table S3. Pairs of second competition experiment | | |  |
| --- | --- | --- | --- |
| Focal cultivars | Neighboring treatments |  |  |
|  | Solitary | Same cultivar | Different cultivars |
| N61 | N61 (n = 30) | N61 vs N61 (n = 30) | N61 vs NBB (n = 8) |
|  |  |  | N61 vs CS (n = 8) |
|  |  |  | N61 vs NP (n = 8) |
|  |  |  | N61 vs *Triticum turgidum* L. subsp. *durum* (Desf.) Husnot cv. ‘Langdon’ (n = 8) |
| NBB | NBB (n = 30) | NBB vs NBB (n = 30) | NBB vs N61 (n = 8) |
|  |  |  | NBB vs CS (n = 8) |
|  |  |  | NBB vs NP (n = 8) |
|  |  |  | NBB vs *T. turgidum* L. subsp. *durum* (Desf.) Husnot cv. ‘Langdon’ (n = 8) |
| CS | CS (n = 30) | CS vs CS (n = 30) | CS vs N61 (n = 8) |
|  |  |  | CS vs NBB (n = 8) |
|  |  |  | CS vs NP (n = 8) |
|  |  |  | CS vs *T. turgidum* L. subsp. *durum* (Desf.) Husnot cv. ‘Langdon’ (n = 8) |
| NP | NP (n = 30) | NP vs NP (n = 30) | NP vs N61 (n = 8) |
|  |  |  | NP vs NBB (n = 8) |
|  |  |  | NP vs CS (n = 8) |
|  |  |  | NP vs *T. turgidum* L. subsp. *durum* (Desf.) Husnot cv. ‘Langdon’ (n = 8) |

| Table S4. Results of generalized linear model examining the effect of root biomass, | | | |
| --- | --- | --- | --- |
| cultivar and their interaction on shoot biomass in distilled water treatment | | | |
| Explanatory variables | *d.f* | *F* | *P*-value^†^ |
| Root biomass | 1 | 49.562 | **< 0.001** |
| Cultivar | 3 | 3.380 | **0.022** |
| N61 vs NBB | 1 | 1.837 | 0.273 |
| N61 vs CS | 1 | 1.489 | 0.275 |
| N61 vs NP | 1 | 0.324 | 0.572 |
| NBB vs CS | 1 | 9.091 | **0.025** |
| NBB vs NP | 1 | 1.899 | 0.273 |
| CS vs NP | 1 | 2.176 | 0.273 |
| Root biomass × Cultivar | 3 | 2.672 | **0.052** |
| N61 vs NBB | 1 | 1.722 | 0.294 |
| N61 vs CS | 1 | 2.257 | 0.294 |
| N61 vs NP | 1 | 7.934 | **0.043** |
| NBB vs CS | 1 | 0.168 | 0.684 |
| NBB vs NP | 1 | 2.083 | 0.294 |
| CS vs NP | 1 | 0.670 | 0.501 |
| Bold indicates *P* < 0.05. |  |  |  |
| ^†^*P*-values were corrected by using the false discovery rate. | | |  |

| Table S5. Results of generalized linear model examining the effect of root biomass, | | | |
| --- | --- | --- | --- |
| treatment and their interaction on shoot biomass in Norin 61 | |  |  |
| Explanatory variables | *d.f* | *F* | *P*-value^†^ |
| Root biomass | 1 | 22.069 | **< 0.001** |
| Treatment | 4 | 6.856 | **< 0.001** |
| DW vs N61 | 1 | 5.838 | **0.028** |
| DW vs NBB | 1 | 7.953 | **0.014** |
| DW vs CS | 1 | 9.177 | **0.010** |
| DW vs NP | 1 | 22.176 | **< 0.001** |
| N61 vs NBB | 1 | 0.158 | 0.693 |
| N61 vs CS | 1 | 0.789 | 0.474 |
| N61 vs NP | 1 | 7.129 | **0.018** |
| NBB vs CS | 1 | 0.486 | 0.544 |
| NBB vs NP | 1 | 9.963 | **0.009** |
| CS vs NP | 1 | 10.515 | **0.009** |
| Root biomass × Treatment | 4 | 4.108 | **0.004** |
| DW vs N61 | 1 | 0.025 | 0.876 |
| DW vs NBB | 1 | 1.500 | 0.284 |
| DW vs CS | 1 | 6.727 | **0.040** |
| DW vs NP | 1 | 11.308 | **0.011** |
| N61 vs NBB | 1 | 1.509 | 0.284 |
| N61 vs CS | 1 | 6.269 | **0.040** |
| N61 vs NP | 1 | 10.678 | **0.011** |
| NBB vs CS | 1 | 2.013 | 0.272 |
| NBB vs NP | 1 | 4.526 | 0.078 |
| CS vs NP | 1 | 0.356 | 0.615 |
| Bold indicates *P* < 0.05. |  |  |  |
| ^†^*P*-values were corrected by using the false discovery rate. | | |  |

| Table S6. Results of analysis by generalized liner model, with each yield parameter as the responsible variables, | | | |
| --- | --- | --- | --- |
| cultivar, treatment and their interaction as the explanatory variables. | | |  |
| Explanatory variables | Responsible variables | Chisq* | *P*-value |
| Total grain biomass per individual (g) | Cultivar | 49.857 | **< 0.001** |
|  | Treatment | 19.248 | **< 0.001** |
|  | Cultivar × Treatment | 9.125 | 0.167 |
| The number of grains | Cultivar | 478.339 | **< 0.001** |
|  | Treatment | 47.961 | **< 0.001** |
|  | Cultivar × Treatment | 97.463 | **< 0.001** |
| Panicle per plant | Cultivar | 10.362 | **0.016** |
|  | Treatment | 3.564 | 0.168 |
|  | Cultivar × Treatment | 20.048 | **0.003** |
| Total grains per panicle (g) | Cultivar | 41.239 | **< 0.001** |
|  | Treatment | 8.983 | **0.011** |
|  | Cultivar × Treatment | 5.460 | 0.486 |
| Thousand-grain weight (g) | Cultivar | 23.216 | **< 0.001** |
|  | Treatment | 11.960 | **0.003** |
|  | Cultivar × Treatment | 5.680 | 0.460 |
| Total aboveground biomass (g) | Cultivar | 47.469 | **< 0.001** |
|  | Treatment | 16.762 | **< 0.001** |
|  | Cultivar × Treatment | 8.477 | 0.205 |
| Harvest index | Cultivar | 17.239 | **< 0.001** |
|  | Treatment | 5.627 | **0.004** |
|  | Cultivar × Treatment | 1.980 | 0.067 |
| Estimated yields (kg/ha) | Cultivar | 49.857 | **< 0.001** |
|  | Treatment | 19.248 | **< 0.001** |
|  | Cultivar × Treatment | 9.125 | 0.167 |
| *In harvest index, the F value is shown instead of the Chisq value. | | | |
| Bold denote significant differences (*P* < 0.05). | |  |  |

| Table S7. Results except for heterospecific, *Triticum turgidum* L. subsp. *durum* (Desf.) Husnot cv. ‘Langdon’ of analysis | | | |
| --- | --- | --- | --- |
| by generalized liner model, with each yield parameter as the responsible variables, cultivar, treatment and their interaction as the explanatory variables. | | | |
| Explanatory variables | Responsible variables | Chisq* | *P*-value |
| Total grain biomass per individual | Cultivar | 54.714 | **< 0.001** |
|  | Treatment | 10.591 | **0.005** |
|  | Cultivar × Treatment | 10.709 | 0.098 |
| The number of grains | Cultivar | 474.272 | **< 0.001** |
|  | Treatment | 36.608 | **< 0.001** |
|  | Cultivar × Treatment | 109.946 | **< 0.001** |
| Panicle per plant | Cultivar | 9.542 | **0.022** |
|  | Treatment | 2.242 | 0.326 |
|  | Cultivar × Treatment | 23.424 | **< 0.001** |
| Total grains per panicle | Cultivar | 69.438 | **< 0.001** |
|  | Treatment | 7.938 | **0.019** |
|  | Cultivar × Treatment | 7.580 | 0.271 |
| Thousand-grain weight (g) | Cultivar | 25.999 | **< 0.001** |
|  | Treatment | 6.225 | **0.044** |
|  | Cultivar × Treatment | 5.531 | 0.478 |
| Total aboveground biomass (g) | Cultivar | 49.595 | **< 0.001** |
|  | Treatment | 7.067 | **0.029** |
|  | Cultivar × Treatment | 10.690 | 0.098 |
| Harvest index | Cultivar | 18.528 | **< 0.001** |
|  | Treatment | 4.986 | **0.007** |
|  | Cultivar × Treatment | 1.165 | 0.324 |
| Estimated yields (kg/ha) | Cultivar | 54.717 | **< 0.001** |
|  | Treatment | 10.592 | **0.005** |
|  | Cultivar × Treatment | 10.709 | 0.098 |
| *In the harvest index, the F value is shown instead of the Chisq value. | | |  |
| Bold denote significant differences (*P* < 0.05). | |  |  |

| Table S8. Yield parameters of each focal cultivar across neighboring treatments | | | | |  |  |  |
| --- | --- | --- | --- | --- | --- | --- | --- |
| Focal cultivars | Treatment | Panicle per plant | Total grains per panicle (g) | Thousand-grain weight (g) | Total aboveground biomass (g) | Harvest index | Estimated yields (kg/ha)* |
| N61 | Solitary | 2.17 ± 0.26^a^ | 21.48 ± 2.46^a^ | 30.43 ± 3.48^a^ | 5.81 ± 0.70^a^ | 0.33 ± 0.04^a^ | 2962.30 ± 361.92^a^ |
|  | Same | 1.85 ± 0.18^a^ | 20.94 ± 1.97^a^ | 32.66 ± 2.90^a^ | 5.58 ± 0.56^a^ | 0.31 ± 0.03^a^ | 2783.58 ± 292.17^a^ |
|  | Different | 1.07 ± 0.31^a^ | 9.35 ± 2.47^b^ | 14.67 ± 3.84^a^ | 3.57 ± 1.59^a^ | 0.15 ± 0.04^b^ | 1260.54 ± 353.54^b^ |
| NBB | Solitary | 3.03 ± 0.30^a^ | 22.28 ± 2.06^a^ | 23.65 ± 2.06^a^ | 5.91 ± 0.54^a^ | 0.31 ± 0.03^a^ | 2511.96 ± 235.16^a^ |
|  | Same | 2.77 ± 0.23^a^ | 20.68 ± 1.57^a^ | 21.54 ± 1.57^a^ | 5.28 ± 0.45^a^ | 0.30 ± 0.02^a^ | 2252.14 ± 202.36^a^ |
|  | Different | 2.04 ± 0.37^a^ | 15.25 ± 2.50^a^ | 14.40 ± 2.32^a^ | 2.72 ± 0.53^b^ | 0.29 ± 0.05^a^ | 1330.30 ± 240.85^b^ |
| CS | Solitary | 2.13 ± 0.25^a^ | 19.15 ± 2.27^a^ | 23.29 ± 2.65^a^ | 4.99 ± 0.58^a^ | 0.26 ± 0.03^a^ | 1972.94 ± 233.78^a^ |
|  | Same | 1.72 ± 0.22^a^ | 14.94 ± 1.87^a^ | 20.90 ± 2.54^a^ | 3.76 ± 0.48^a^ | 0.23 ± 0.03^a^ | 1606.34 ± 203.26^a^ |
|  | Different | 1.77 ± 0.32^a^ | 14.03 ± 2.44^a^ | 16.56 ± 2.72^a^ | 3.36 ± 0.67^a^ | 0.23 ± 0.04^a^ | 1470.73 ± 291.47^a^ |
| NP | Solitary | 2.07 ± 0.38^a^ | 8.42 ± 1.54^a^ | 19.18 ± 3.42^a^ | 2.84 ± 0.53^a^ | 0.21 ± 0.04^a^ | 1295.78 ± 244.11^a^ |
|  | Same | 0.95 ± 0.23^a^ | 3.54 ± 0.85^b^ | 9.11 ± 2.17^a^ | 1.25 ± 0.33^b^ | 0.10 ± 0.02^b^ | 582.97 ± 159.78^b^ |
|  | Different | 0.39 ± 0.18^b^ | 3.00 ± 1.17^b^ | 6.01 ± 2.28^a^ | 0.44 ± 0.21^b^ | 0.09 ± 0.04^b^ | 219.30 ± 104.50^b^ |
| *This value was estimated from the results of grain biomass per individual and pot size. | | | | |  |  |  |
| Different letters denote significant differences (*P* < 0.05). | | | |  |  |  |  |

4 mL

Distilled water

Same cultivar exudate

Different cultivar exudate

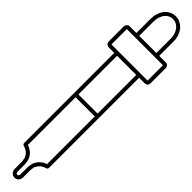

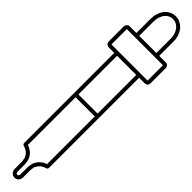

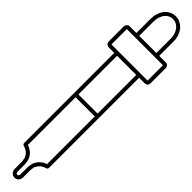

**（a）**

**（b）**

Silica sand

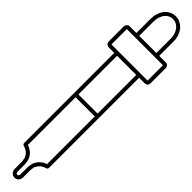

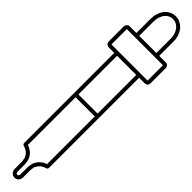

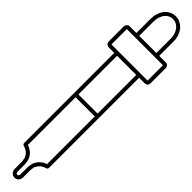

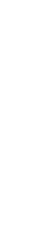

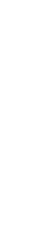

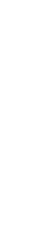

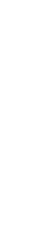

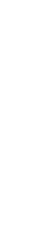

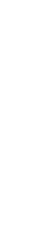

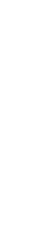

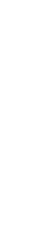

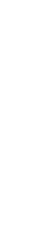

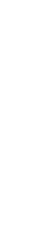

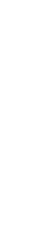

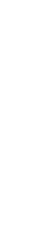

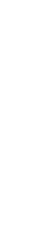

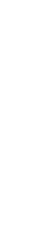

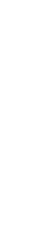

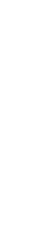

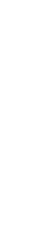

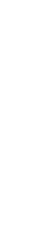

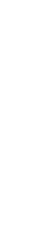

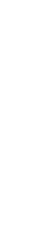

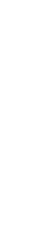

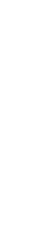

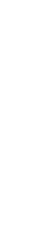

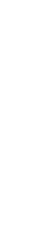

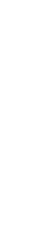

Fig.S1 Methods of root exudate collection and application. **(a)** Root exudate collection: fifty seedlings of each common wheat *Triticum aestivum* L. cultivar were grown under hydroponic conditions with 300mL distilled water. **(b)** Root exudate application: 4 mL distilled water or the root exudates of the same cultivar or different cultivars were applied to seedlings of each cultivar transplanted into a 10 mL pipette tip containing silica sand every day.

ALT TEXT: Figures showed (a) root exudate collection and (b) application 4mL of root exudate.

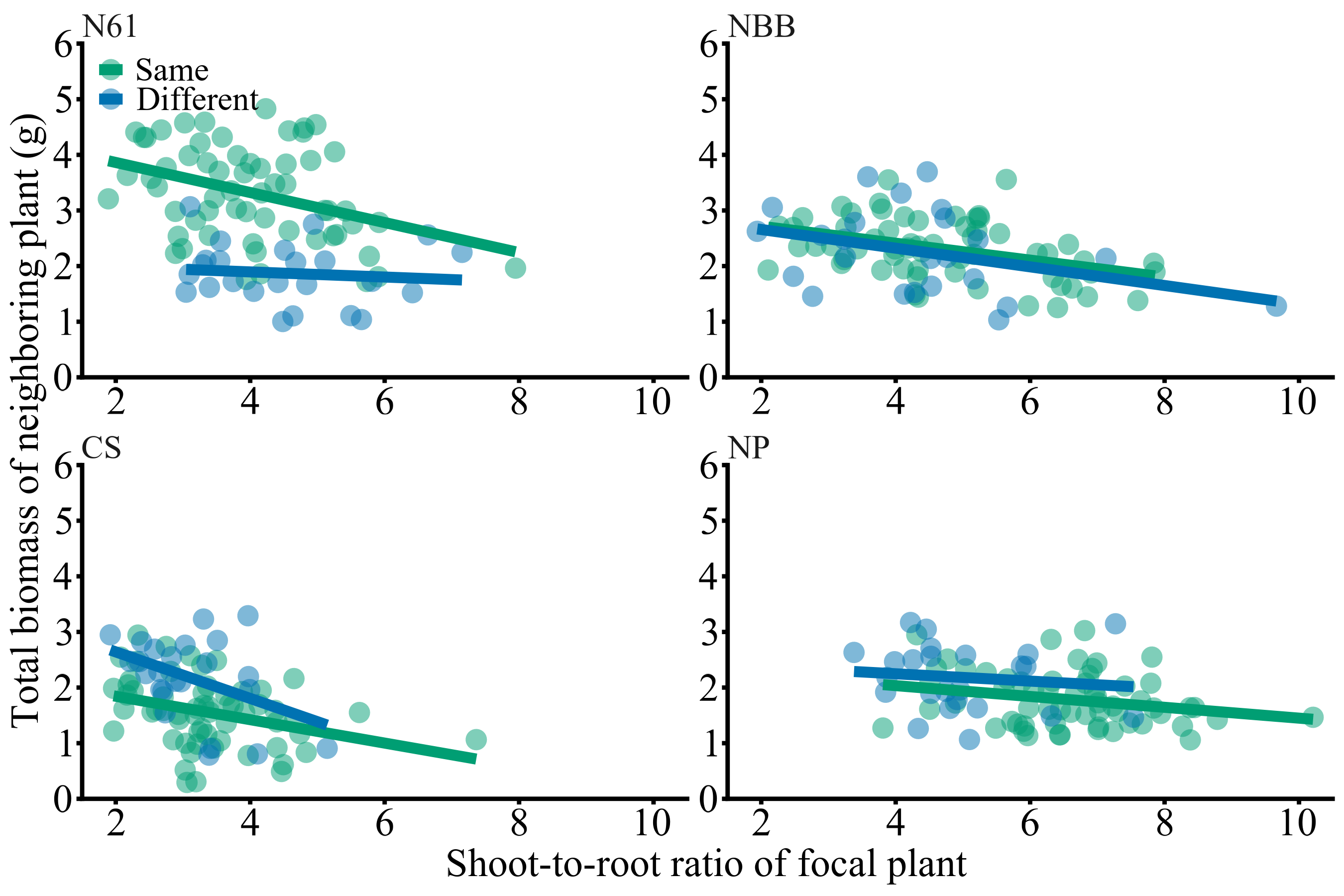

Fig.S2 Relationship between shoot-to-root allocation and competitor biomass of each common wheat cultivar at 38 days. Each line indicates significant correlations (LMM, *P* < 0.001).

ALT TEXT: This figure showed effects of shoot-to-root ratio of focal wheat plants on total biomass of neighboring plants for neighboring treatments.

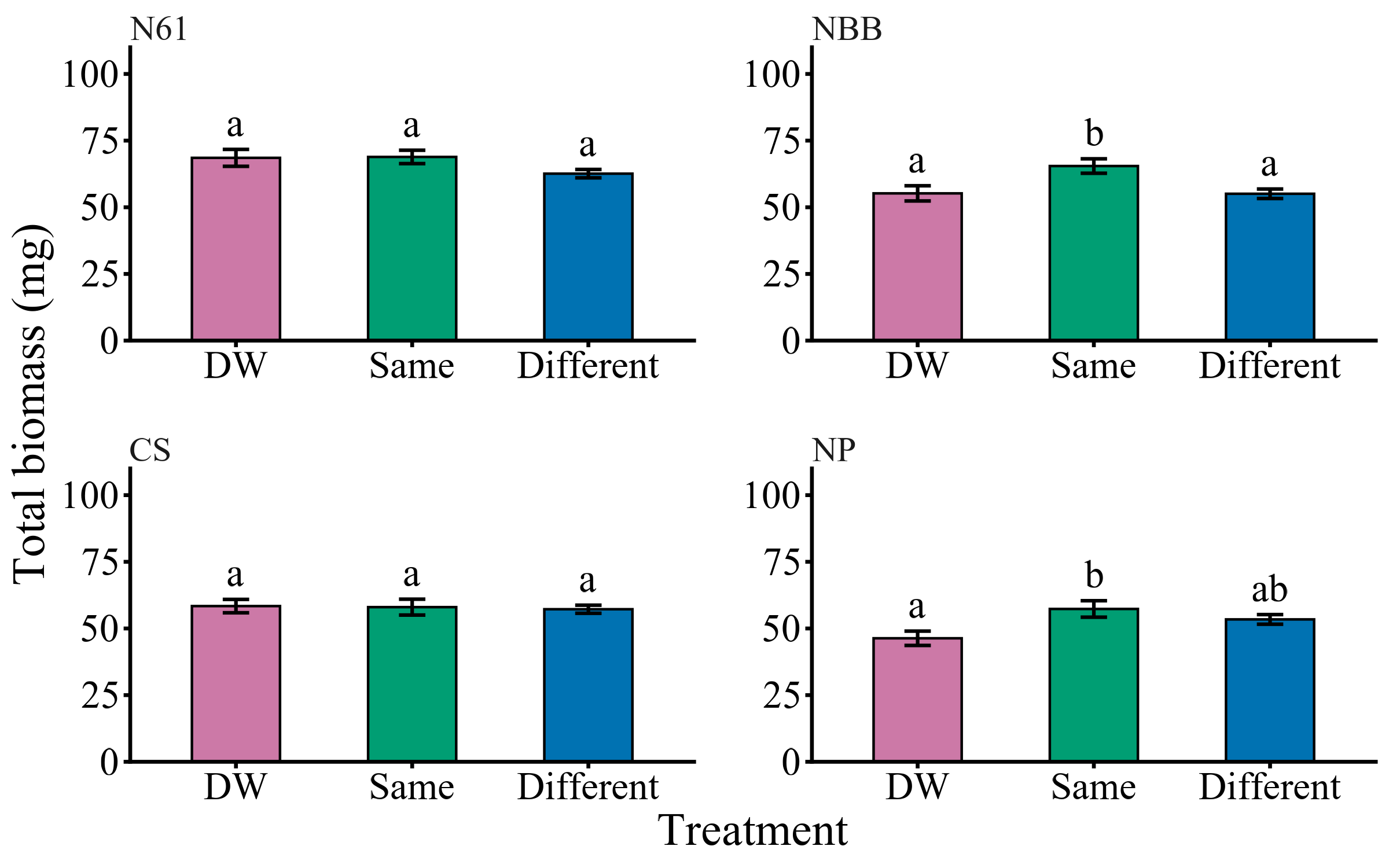

Fig.S3 Total biomass of each common wheat *Triticum aestivum L.* cultivar in each experimental condition: exposed to distilled water (DW), the root exudates from the same cultivar (Same) or different cultivars (Different). N61, NBB, CS and NP stand for “Norin 61”, “Nobeokabouzu”, “Chinese Spring” and “KU-7113”, respectively. Bars represent SE. Different letters show significant differences (GLM, *P* < 0.05).

ALT TEXT: Graphs showed total biomass of each cultivar depending on root exudate treatments.
